# Modularity in the DCC extracellular domain elicits distinct effects on axon guidance

**DOI:** 10.64898/2025.12.11.693717

**Authors:** Evelyn C. Avilés, Zachary DeLoughery, Andrea Yung, Jia-Huai Wang, Rob Meijers, Lisa Goodrich

## Abstract

Complex neuronal circuits arise from a small set of cell-surface receptors that position neurons, promote axon extension, and define synaptic connections. A central receptor is Deleted in Colon Cancer (DCC), which mediates both short- and long-range axon guidance and confines migrating neurons to the central nervous system. DCC’s versatility reflects its ability to interact at distinct sites of its extracellular domain with two ligands, Netrin-1 and Draxin, which also bind to each other. Alternative splicing further alters the Netrin-1 binding site and modulates affinity. By generating two mouse lines with mutations that selectively impair DCC binding to Netrin-1 and/or Draxin, we show that molecular modularity within the DCC extracellular domain is essential for precise circuit assembly. An eight–amino acid insertion in the DCC^long^ isoform is required for Netrin-1–dependent long-range commissural axon guidance in the spinal cord. Conversely, isoleucine 372 in the Draxin binding site enables DCC clustering and is necessary for all known DCC functions, including axon guidance in the spinal cord and retina and neuronal migration in the brainstem. Draxin also supports long-range commissural guidance, but mutations in its binding site cause stronger defects. These results underscore how DCC’s distinct modules drive specific developmental responses.

## Introduction

In the developing nervous system, growing axons respond to positive and negative cues that collectively steer their growth cones along specific paths towards their final targets. Although initial studies often assigned molecules as chemoattractants or chemorepellants, it is increasingly apparent that individual molecules can elicit a range of outcomes depending on the context in which they are received (Chédotal 2019; Comer et al. 2019; Kolodkin and Tessier-Lavigne 2011). An important example is the classic chemoattractant Netrin-1. Originally characterized for its ability to attract and turn axons across a long range, Netrin-1 can also play a more permissive role that depends on short range interactions (Varadarajan et al. 2017; Dominici et al. 2017; Hand and Kolodkin 2017). Consistent with the importance of short-range permissive effects, broad overexpression of Netrin-1 by Nestin+ precursors is sufficient to rescue pontine motor neuron migration defects in the hindbrain (Yung et al. 2018). Both long- and short-range activities have also been described in the retina (Vigouroux et al. 2020; Deiner et al. 1997). By investigating how Netrin-1 elicits distinct effects on the same axons in different contexts, we can better understand how molecular versatility contributes to reliable wiring of the nervous system.

One way that Netrin-1 activities can be tuned is through its receptor Deleted in Colorectal Cancer (DCC), an Ig superfamily protein. Although there is a minor contribution from a related receptor, Neogenin, most of Netrin-1’s effects on commissural axons appear to be through DCC (Kolodkin and Tessier-Lavigne 2011) (**Figure 1A**). One potential source of signaling variability is the modularity of its extracellular domain (ECD). Alternative splicing of DCC generates short (*Dcc*^short^) or long (*Dcc*^long^) isoforms (Cooper et al. 1995). *Dcc*^short^ lacks a 60bp stretch of exon 17 that encodes a 20 amino acids linker sequence located between fibronectin type III domains (FN) FN4 and FN5 in the extracellular domain of DCC^long^ (Cooper et al. 1995; Dailey-Krempel et al. 2023; Finci et al. 2017) (**Figure 1B**). Due to alternative splicing, the FN4-5 linker varies from 8 residues in DCC^short^ to 28 residues in DCC^long^. Based on crystallographic structural studies, the length of the linker is predicted to determine the availability of all three Netrin-1 binding sites (EGF3, EGF1/2, and LN) to interact with each molecule of DCC and could therefore modulate the strength of DCC/Netrin-1 binding (Finci et al. 2017). Binding to all three sites in DCC^long^ may also provide higher stability to the FN5 domain and promote oligomerization and clustering of the Netrin-1/DCC pair to promote axon attraction (Finci et al. 2015, 2017, 2014). By contrast, the lack of this linker region makes it more likely that Netrin-1 binds to sites on two DCC^short^ molecules, thereby creating a molecular bridge. Cell surface expression and ELISA assays showed that netrin binding is affected by the linker length between domains FN4 and FN5 (Finci et al 2017), and heparan sulfate binding affected Netrin-1/DCC interactions differently for the short and long DCC isoforms (Priest et al. 2024). The short isoform is highly expressed in embryos and this shifts to exclusive expression of the long isoform as development proceeds (Cooper et al. 1995; Leggere et al. 2016), but it is unclear how this change in expression impacts Netrin-1 signaling output. One possibility is that DCC^long^ mediates long range guidance, as DCC^long^ and not DCC^short^ was able to rescue commissural guidance in *Nova* mutant mouse embryos (Leggere et al. 2016).

**Figure 1:**
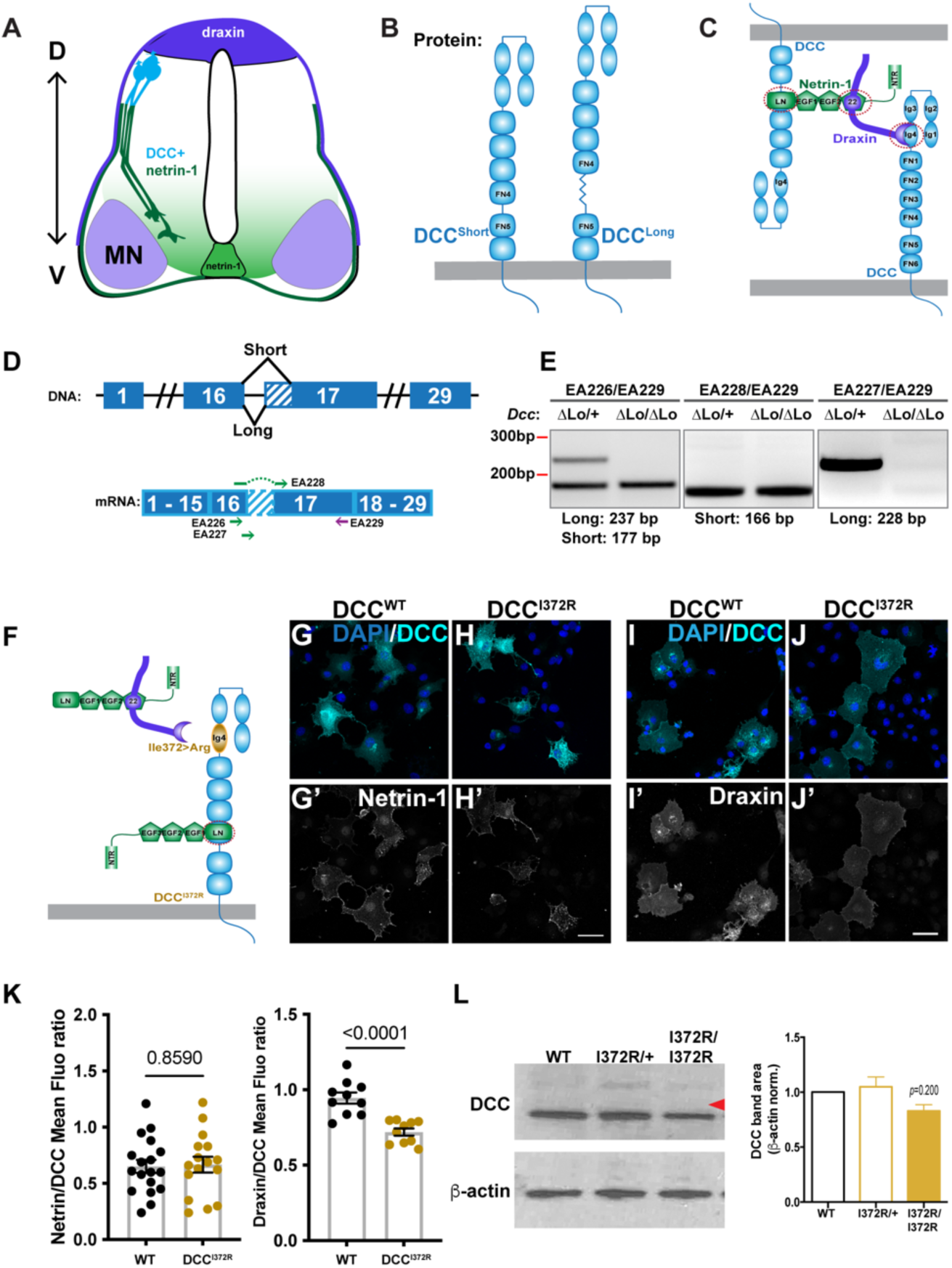
*Dcc*^I372R^ and *Dcc*^Δlong^ mutant variants. **(A)** Representation of a developing spinal cord cross-section expressing Draxin, Netrin-1 and DCC in commissural axons. MN: motor neurons; D: dorsal; V: ventral. **(B)** Representation of the protein products of alternative splicing of the *Dcc* gene. **(C)** Representation of the intercellular tripartite complex between DCC, Netrin-1 and Draxin. **(D)** Representation of the genomic locus of the *Dcc* gene showing the alternative splicing event around exon 17 (upper panel), and the alignment of the primers to detect the short, the long or both isoforms (lower panel). **(E)** RT-PCR to detect isoform-specific fragments of the *Dcc* gene confirming the specific *Dcc*^Δlong^ mutation in the mouse line. **(F)** Representation of the *Dcc*^I372R^ mutation predicted to interfere with Draxin binding. **(G,H)** Overexpression of a plasmid coding for DCC^WT^ (G) or DCC^I372R^ (H) in Cos7 cells stained for DCC and for Netrin-1, which was added to the culture. **(I,J)** Overexpression of a plasmid coding for DCC^WT^ (I) or DCC^I372R^ (J) in Cos7 cells stained for DCC and for Draxin, which was added to the culture. **(K)** Quantification of Netrin-1 mean fluorescence on DCC+ cells. DCC^WT^: 0.65 ± 0.06, n=18 ROI; DCC^I372R^: 0.67 ± 0.07, n=16 ROI; both conditions in quadruplicate; Mann-Whitney test (left). Quantification of Draxin mean fluorescence on DCC+ cells. DCC^WT^: 0.95 ± 0.04, n=10 ROI; DCC^I372R^: 0.72 ± 0.02, n=10 ROI; both conditions in quadruplicate, t test (right). **(L)** Western blot to DCC shows full length expression of the DCC^I372R^ protein. Red arrowhead indicates 150 kDa. Scale bars: 50µm.

Netrin-1/DCC activity may be further modulated by interactions with other ligands, such as Draxin (Ahmed et al. 2011; Liu et al. 2018; Meijers et al. 2020). Draxin is a guidance molecule involved in commissure formation in the brain (Shinmyo 2025; Islam et al. 2009; Ahmed and Shinmyo 2021). Based on structural data, it has been proposed that a tripartite complex between DCC, Netrin-1, and Draxin mediates axon fasciculation by forming a bridge between DCC molecules on apposing surfaces (Liu et al. 2018; Meijers et al. 2020) (**Figure 1C**). Despite Draxin’s ability to bind strongly to both Netrin-1 and DCC, *Draxin* mutants do not phenocopy either *Ntn1* or *Dcc* null mutants, but instead are fertile and viable, with abnormal forebrain commissures and subtle defects in commissural axon guidance in the spinal cord (Islam et al. 2009). Draxin secreted from the dorsal spinal cord may act as a repulsive guidance cue (Islam et al. 2009), consistent with its ability to repel axons *in vitro* via a mechanism that involves DCC (Ahmed et al. 2011). Therefore, DCC may mediate axonal attraction through Netrin-1 and repulsion through Draxin, respectively. In this model, loss of Draxin-DCC dependent repulsion may be rescued by persistent attraction via Netrin-1. Whether DCC^short^ and/or DCC^long^ is involved in Draxin-dependent activities is not known.

Here, we sought to dissect DCC’s versatile effects on developing neurons by selectively ablating DCC^long^ and by altering the Draxin binding site. Comparison between the effects of each mutation and the loss of Draxin altogether suggests that these changes to the DCC-ECD lead to fundamentally different effects that may involve ligands other than Draxin.

## Results

### Molecular dissection of DCC signaling functions

Previous work raises the possibility that DCC^long^ mediates long distance guidance, whereas DCC^short^ and Draxin-bound DCC are involved in adhesion and fasciculation (Leggere et al., 2016; Liu et al., 2018). In the embryonic spinal cord, Netrin-1 can act both adhesively, for instance by mediating fasciculation of DCC+ axons, and instructively, by inducing DCC+ axons to turn towards the floor plate (Meijers et al. 2020). On the other hand, Draxin is proposed to serve as a repulsive cue that guides axons to the ventral-medial areas (Islam et al. 2009) (**Figure 1A-C**), though how this affects Netrin1-DCC signaling outcomes is not known.

To test how these different modes of DCC signaling impact neuronal development, we used CRISPR/Cas-9 technology to selectively disrupt signaling mediated by DCC^long^ and Draxin-bound DCC. To eliminate the long isoform, we targeted the 5’ terminus of exon 17, which contains an internal splice acceptor sequence used to produce DCC^short^. In *Dcc*^Δlong^ mice, 25 nucleotides immediately upstream of exon 17 were deleted, thereby preventing splicing at the 5’-end of the exon while maintaining splicing to the internal splice acceptor to produce *Dcc*^short^ (**Figure 1D**). RT-PCR of *Dcc*^Δlong/Δlong^ tissue confirmed loss of *Dcc*^long^ expression (**Figure 1E**). Therefore, *Dcc*^Δlong/Δlong^ mice only produce DCC^short^.

To investigate the impact of Draxin binding (**Figure 1C**), we created a mouse strain where we changed the isoleucine at position 372 into an arginine (*Dcc*^I372R^ line), as the introduction of a basic residue in this position is strongly predicted to prevent Draxin-DCC binding without affecting Netrin-1-DCC binding (Liu et al. 2018) (**Figure 1F**). Indeed, soluble Netrin-1 accumulates on the surface of DCC^I372R^-expressing cells, but Draxin binding is significantly reduced (**Figure 1G-K**). In mice homozygous for this mutation (*Dcc*^I372R/I372^), neither the size nor the overall amount of DCC changed relative to controls (**Figure 1L**). Thus, these two new *Dcc* alleles affect different features of the DCC ECD predicted to mediate binding of Netrin-1 or Draxin.

Breeding of the *Dcc*^Δlong^ and *Dcc*^I372R^ strains revealed clear differences in contributions of each form of DCC to mouse development. Whereas *Dcc* null mutants die at birth, *Dcc*^Δlong^ homozygotes were viable and fertile, with no obvious morphological or behavioral defects. By contrast, although *Dcc*^I372R^ homozygotes were generated at the expected Mendelian ratio prior to birth (64/258 embryos, 24.8%), few *Dcc*^I372R^ homozygotes survived the first day of life (12 out of 96 animals born to heterozygous crosses, 12.5%). These survivors were much smaller than their wild-type and heterozygous littermates and most of them (7 out of 12 animals) died perinatally; although this number might be underestimated as perinatal death is often followed by maternal cannibalism. Only 5 out of 12 *Dcc*^I372R^ homozygous postnatal animals (41.7%) reached 4 weeks of age.

### *Dcc*_short_ and *Dcc*_long_ are differentially expressed in the developing spinal cord

To elucidate how diverse forms of DCC signaling influence axon guidance, we focused on the guidance of commissural axons in the embryonic spinal cord. Analysis of *Dcc_short_* and *Dcc_long_* expression patterns revealed that both isoforms are present in developing commissural neurons, with expression of *Dcc*_long_ also more ventrally. Using isoform-specific probes (**Figure 2A,B**) and BaseScope *in situ* hybridization technology (Baker et al. 2017; Avilés et al. 2022), we found that *Dcc*_short_ was present at high levels in the E11.5 dorsal spinal cord, where commissural cell bodies reside (**Figure 2C**). *Dcc*_long_ was also expressed in the dorsal spinal cord, as well as in the medial ventral area of the spinal cord (**Figure 2E**). In *Dcc*^Δlong/Δlong^ mutants, *Dcc*_short_ expression expanded ventrally (**Figure 2D**). Consistent with the RT-PCR results (**Figure 1E**), no *Dcc*_long_ was seen in BaseScope *in situ* hybridization of *Dcc*^Δlong/Δlong^ mutant spinal cord samples (**Figure 2F**). We were unable to differentiate DCC isoforms at the protein level due to the lack of isoform-specific antibodies. Nonetheless, immunostaining confirmed that DCC protein, which likely corresponds to DCC^short^, is still present and normally localized to subsets of neurons in the embryonic spinal cord of *Dcc*^Δlong/Δlong^ mutants (**Figure S1A,B**).

**Figure 2:**
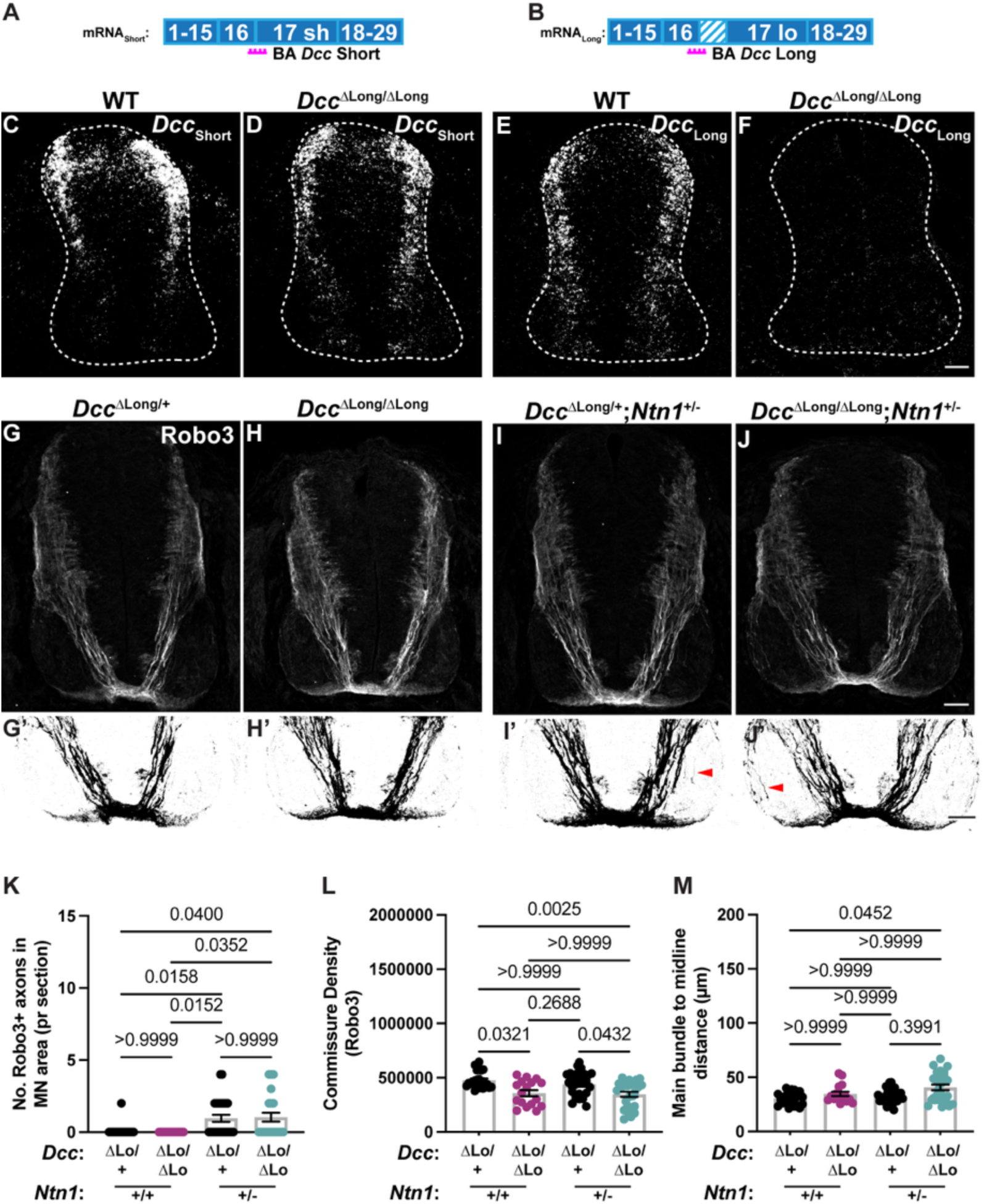
DCC^long^ cooperates with Netrin-1 to promote commissural axon guidance to the floor plate. **(A,B)** Representation of the exonic splicing in the short (A) and long (B) mRNA form of the *Dcc* gene. The Basescope (BA) probes to the short and long isoforms are mapped in magenta. **(C,D)** Basescope *in situ* hybridization to the *Dcc*_short_ in wild type (C) and *Dcc*^Δlong/Δlong^ (D) E11.5 spinal cords. **(E,F)** Basescope *in situ* hybridization to the *Dcc*_long_ in wild type (E) and *Dcc*^Δlong/Δlong^ (F) E11.5 spinal cords. **(G-J)** Robo3 staining of *Dcc*^Δlong/+^ (G), *Dcc*^Δlong/Δlong^ (H), *Dcc*^Δlong/+^;*Ntn1*^+/-^ (I), and *Dcc*^Δlong/Δlong^;*Ntn1*^+/-^ (J) E11.5 spinal cords. The most ventral area of the spinal cords is shown in binarized images in G’-J’. **(K)** Quantification of the number of Robo3+ axons in the motor neuron (MN) area. *Dcc*^Δlong/+^: 0.08 ± 0.08, n=24 sections from 6 mice; *Dcc*^Δlong/Δlong^: 0.00 ± 0.00, n=17 sections from 4 mice; *Dcc*^Δlong/+^;*Ntn1*^+/-^: 0.96 ± 0.24, n=27 sections from 7 mice; *Dcc*^Δlong/Δlong^;*Ntn1*^+/-^: 1.04 ± 0.31, n=26 sections from 7 mice. Kruskal-Wallis test. **(L)** Quantification of the commissure density by Robo3 staining. *Dcc*^Δlong/+^: 477489 ± 15158, n=24 sections from 6 mice; *Dcc*^Δlong/Δlong^: 357951 ± 28155, n=17 sections from 4 mice; *Dcc*^Δlong/+^;*Ntn1*^+/-^: 438363 ± 206648, n=28 sections from 7 mice; *Dcc*^Δlong/Δlong^;*Ntn1*^+/-^: 347356 ± 23556, n=26 sections from 7 mice. Kruskal-Wallis test. **(M)** Quantification of the distance between main bundle to the midline (µm). *Dcc*^Δlong/+^: 30.64 ± 1.18, n=24 sections from 6 mice; *Dcc*^Δlong/Δlong^: 34.74 ± 2.00, n=17 sections from 4 mice; *Dcc*^Δlong/+^;*Ntn1*^+/-^: 32.70 ± 1.25, n=28 sections from 7 mice; *Dcc*^Δlong/Δlong^;*Ntn1*^+/-^: 40.67 ± 2.63, n=26 sections from 7 mice. Kruskal-Wallis test. Scale bars: 50µm.

### Distinct DCC-ECD modules guide commissural axons to the floor plate

Unlike *Dcc* null mice, commissural axon trajectories appeared largely normal in *Dcc*^Δlong/Δlong^ mutants. Robo3+ commissural axons in *Dcc*^Δlong/Δlong^ mutants followed normal trajectories through the embryonic spinal cord (**Figure 2G,H,K)**, with no evidence for guidance defects seen in *Dcc*^-/-^ mice (Fazeli et al. 1997; Varadarajan et al. 2017), such as stray axons growing towards the roof plate or invasion of the motor column (**Figure S2G,H**). However, the commissure was slightly (∼25%) but significantly thinner in E11.5 *Dcc*^Δlong/Δlong^ than in WT embryos (**Figure 2G,H,L, Figure S1C**), consistent with the finding that there is a delayed entrance of axons to the floor plate in a similar mouse strain, DCC_s_, that also expresses only DCC^short^ (Dailey-Krempel et al. 2023). To test whether additional DCC^long^ roles might be uncovered when less Netrin-1 is available, we removed one copy of the Netrin-1 gene (*Ntn1*) from the *Dcc*^Δlong/Δlong^ background. This reduction in *Ntn1* gene dosage led to the appearance of axon guidance defects in the motor columns (**Figure 2I,J,K**) that were similar to what happens in *Ntn1* null homozygotes (**Figure S1F,G**). This effect was seen both in the *Dcc*^Δlong/+^ and *Dcc*^Δlong/Δlong^ background (**Figure 2I,J,K**). To further evaluate aspects of long range Netrin1-dependent guidance, we quantified the lateral displacement of pre-crossing axons in the ventral spinal cord by measuring the distance of the main bundle to the midline (Wu et al. 2019) (**Figure S1D**). As predicted, there was a subtle lateral displacement of Robo3+ axons in *Dcc*^Δlong/Δlong^;*Ntn1*^+/-^ ventral spinal cord (**Figure 2J,M**) that was not present in *Ntn1*^+/-^heterozygous animals (**Figure S1E,G,H**) or in *Dcc*^Δlong/Δlong^ spinal cords (**Figure 2H,M**). Collectively, these subtle phenotypes support the hypothesis that DCC^long^ is required for Netrin-1 long-range attraction and demonstrate that DCC^short^ can substitute for DCC^long^ to guide commissural axons to the midline.

In stark contrast to *Dcc*^Δlong/Δlong^ mutants, commissural axon trajectories were highly abnormal in *Dcc*^I372R/I372R^ animals, with defects reminiscent of those in *Dcc*^-/-^ animals (Fazeli et al. 1997) (**Figure 3A-C**). Robo3+ commissural axons were defasciculated (**Figure 3D**) and often grew inappropriately towards the dorsal spinal cord (**Figure 3E**) and into the motor columns (**Figure 3F**). In addition, the midline commissure was significantly thinner in the *Dcc*^I372R/I372R^ mutant spinal cord (**Figure 3G**), also evident in whole mount open book preparations (**Figure 3H-J)** and by staining for DCC and NF (**Figure S2A-E**). The range and severity of phenotypes was similar to those in *Dcc*^-/-^ mutant spinal cords (**Figure S2F-I**). Therefore, whereas loss of DCC^long^ has only subtle effects consistent with a loss of long-range signaling, altering Draxin’s binding site strongly disrupts DCC’s overall ability to guide commissural axons towards the midline.

**Figure 3:**
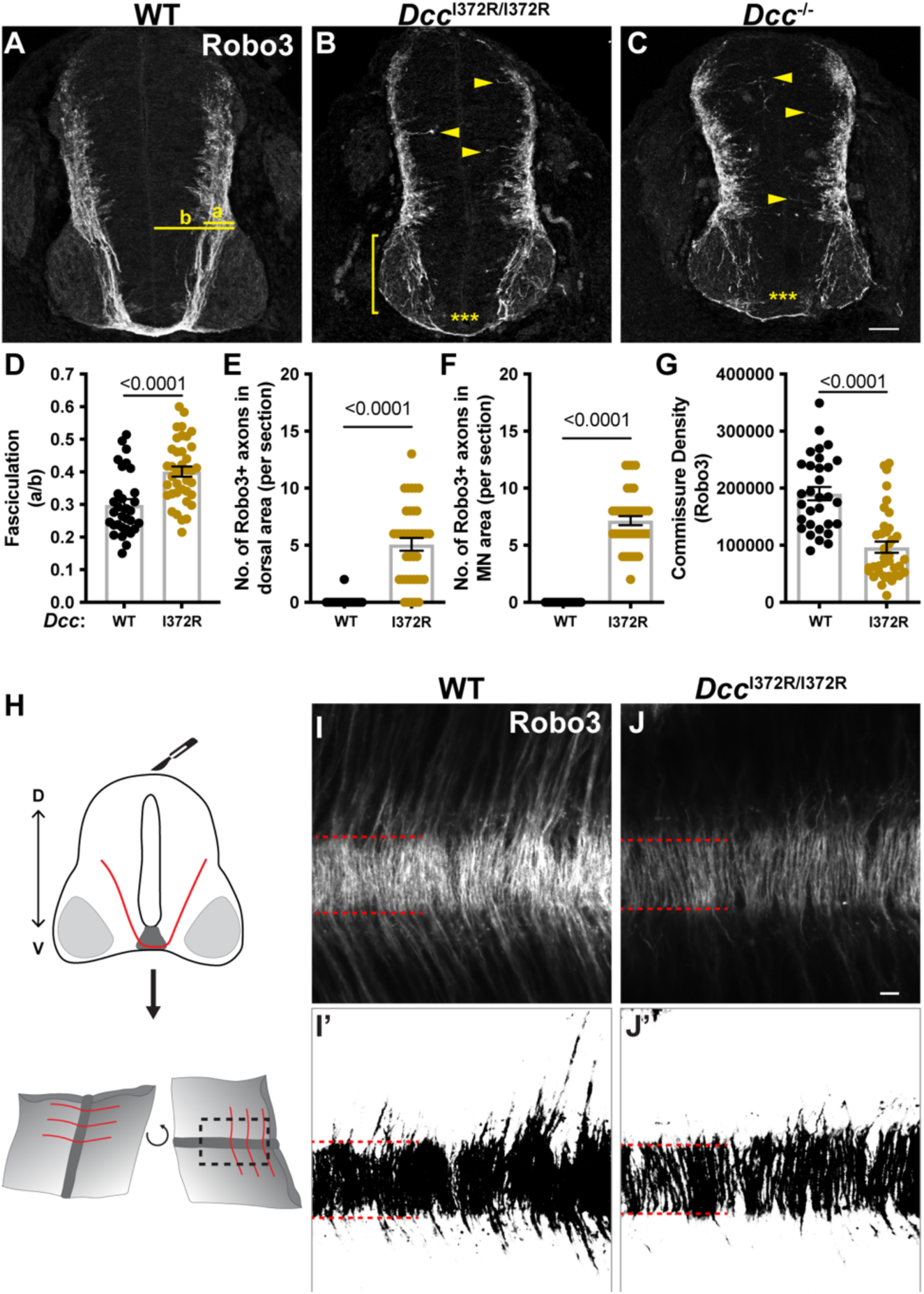
*Dcc*^I372R/I372R^ mutant mice exhibit penetrant commissural axon guidance defects. **(A)** Immunostaining of Robo3 on E11.5 spinal cord cross sections of WT animals. "a" indicates the distance from the pial surface to where the most medial Robo3+ axon is located, and "b" represents the distance from the pial surface to the midline. **(B,C)** Immunostaining of Robo3 on E11.5 spinal cord cross sections of *Dcc*^I372R/I372R^ (B) and *Dcc*^-/-^ (C) animals. **(D)** Quantification of the fasciculation parameter (a/b). WT: 0.30 ± 0.02, n=31; *Dcc*^I372R/I372R^: 0.40 ± 0.09, n=38; *t* test. 8 mice per condition. **(E)** Quantification of the number of Robo3+ axons in the dorsal area. WT: 0.06 ± 0.06, n=31; *Dcc*^I372R/I372R^: 5.08 ± 0.56, n=38; Mann-Whitney test. 8 mice per condition. **(F)** Quantification of the number of Robo3+ axons in the motor neuron (MN) area. WT: 0.00 ± 0.00, n=31; *Dcc*^I372R/I372R^: 7.16 ± 0.40, n=38; Mann-Whitney test. 8 mice per condition. **(G)** Quantification of the commissure density by Robo3 staining. WT: 190569 ± 11763, n=31; *Dcc*^I372R/I372R^: 96582 ± 9866, n=38; Mann-Whitney test. 8 mice per condition. **(H)** Schematic representation of an open book preparation. **(I)** Whole-mount staining for Robo3 on WT spinal cords. **(J)** Whole-mount staining for Robo3 on *Dcc*^I372R/I372R^ spinal cords. I’ and J’ show binarized I and J images. Red lines depict the floor plate. Scale bars: 50µm (C); 20µm (J’).

### Draxin is required for commissural axon guidance near the floor plate but not dorsally

Draxin is a secreted molecule originally characterized as a repulsive cue that acts in the dorsal spinal cord to direct commissural axons ventrally. Draxin binds to Netrin-1 and also to DCC in a site that is distant from the Netrin-1 binding site (Liu et al. 2018). Structurally, this tripartite complex suggests that Draxin brings two DCC molecules together by binding directly to one DCC in one pair and to Netrin-1 in the other (**Figure 1C**). This arrangement is well-suited to enable adhesion and fasciculation, but the precise contributions of Draxin to commissural axon guidance remain unclear.

Analysis of *Draxin*^-/-^ mutant mice (Islam et al. 2009) revealed a role for Draxin in the ventral spinal cord that was not previously described. Both fluorescent *in situ* hybridization and immunostaining confirmed the presence of Draxin in the dorsal spinal cord but also revealed additional sources ventrally (**Figure 4A,C**). This signal was absent in *Draxin*^-/-^ mice (**Figure 4B,D**), confirming both that Draxin is normally produced both dorsally and ventrally and that these sources are eliminated effectively in *Draxin^-/-^* mutants. Despite the complete absence of Draxin, Robo3+ commissural axons followed a stereotypical path from the dorsal to the ventral spinal cord (**Figure 4E-H**), as shown previously (Islam et al. 2009). However, the spinal cord commissure was obviously thinner in *Draxin^-/-^* mice (**Figure 4E’,F’,I**), confirmed by staining for NF and DCC (**Figure S3A-F**), consistent with the thinning of the anterior commissure previously reported (Islam et al. 2009). In addition, Robo3+ axons were displaced laterally and no longer followed a sharp diagonal path to the midline (**Figure 4 F’,J**). This phenotype is similar to what happens when Netrin-1 is eliminated from the floor plate, which has been attributed to Netrin-1’s long-range action (Wu et al. 2019).

**Figure 4:**
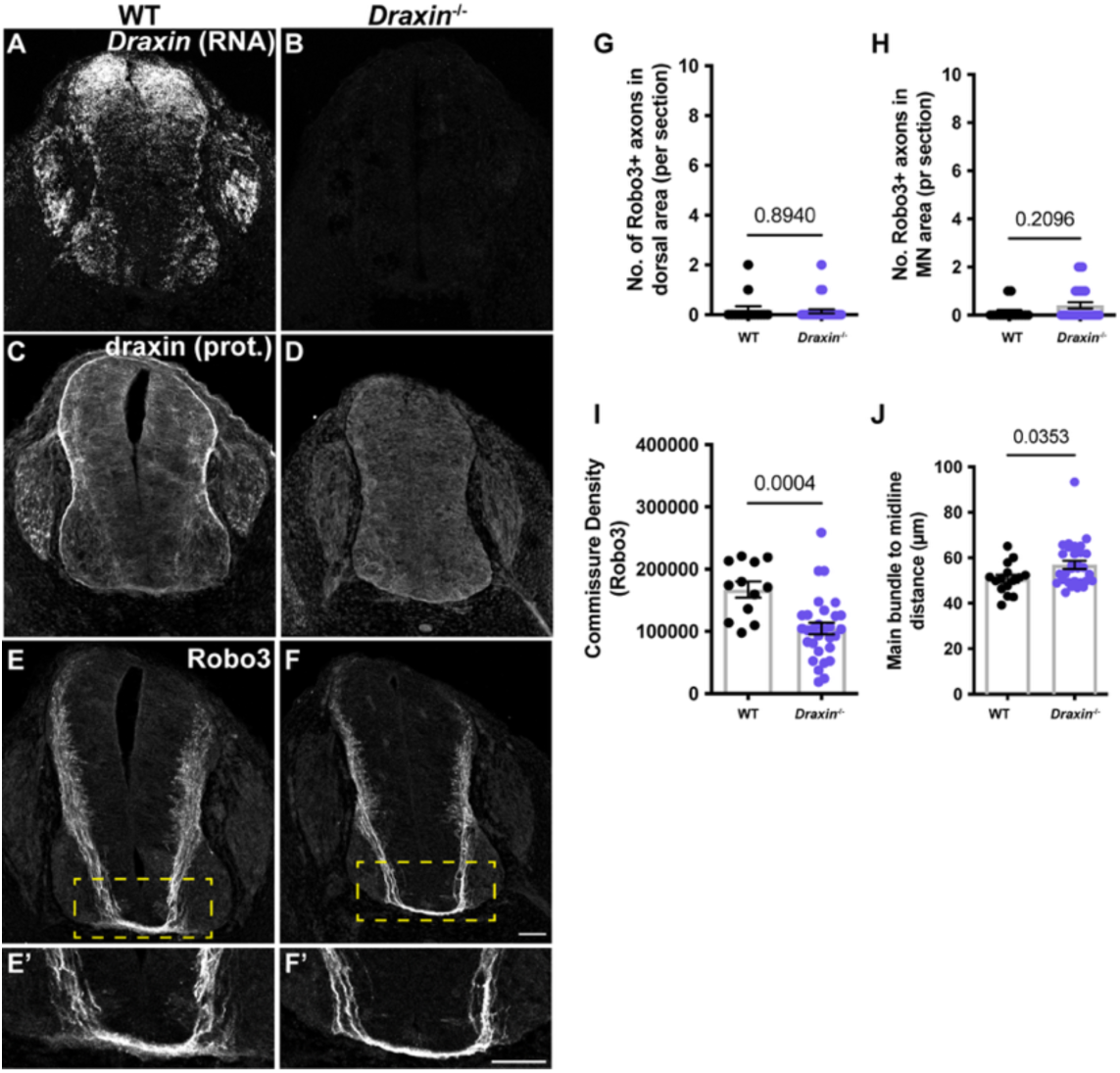
*Draxin* is required for ventral axon guidance in the developing spinal cord. **(A,B)** RNAscope *in situ* hybridization to *Draxin* RNA in wild type (A) and *Draxin*^-/-^ mutant (B) E11.5 spinal cords. **(C,D)** Draxin immunostaining in wild type (C) and *Draxin*^-/-^ mutant (D) E11.5 spinal cords. **(E,F)** Robo3 immunostaining in wild type (E) and *Draxin*^-/-^ mutant (F) E11.5 spinal cords. Regions delineated in the yellow squares are shown at higher magnification in E’ and F’. **(G)** Quantification of the number of Robo3+ axons incorrectly located in the dorsal area of the spinal cord. WT: 0.20 ± 0.14, n=15; *Draxin*^-/-^: 0.14 ± 0.08, n=29. Mann-Whitney test. **(H)** Quantification of the number of Robo3+ axons in the motor neuron (MN) area. WT: 0.13 ± 0.09, n=15; *Draxin*^-/-^: 0.41 ± 0.13, n=29. Mann-Whitney test. **(I)** Quantification of the commissure density by Robo3 staining. WT: 167249 ± 12873, n=12; *Draxin*^-/-^: 104583 ± 9466, n=29. Mann-Whitney test. **(J)** Quantification of the distance between main bundle to the midline (µm). WT: 50.80 ± 1.75, n=15; *Draxin*^-/-^: 56.94 ± 1.79, n=30. Mann-Whitney test. WT: 3 mice, *Draxin*^-/-^: 7 mice. Scale bars: 50µm.

These findings suggest that Draxin has a similar effect on long-range commissural axon guidance as Netrin-1. Additionally, we find that *Draxin* mutant phenotypes are much milder than those observed in *Dcc*^I372R/I372R^ mutants. This raises the possibility either that additional ligands require I372 for binding to the DCC ECD or that the I372R mutation prevents DCC from mediating Netrin-1 signaling in other ways.

### Draxin does not counteract Netrin-1 signaling in a *Dcc*^I372R/I372R^ mutant background

One key difference between the *Dcc*^I372R/I372R^ and *Draxin*^-/-^ strains is the availability of Draxin, which can antagonize Netrin-1 by binding to it (Gao et al. 2015). We hypothesized that Draxin effectively sequesters Netrin-1 due to its inability to bind to DCC^I372R^. If this is true, then loss of Draxin should rescue aspects of the *Dcc*^I372R/I372R^ phenotype. To test this idea, we generated *Draxin*^-/-^;*Dcc*^I372R/I372R^ double mutants and examined them for axon guidance defects.

Contrary to our hypothesis, axon guidance was highly abnormal in the spinal cord of E11.5 *Draxin*^-/-^;*Dcc*^I372R/I372R^ double mutants (**Figure S4A-D**), and none of the analyzed parameters were rescued (**Figure S4E-H**). There was also no change in DCC expression, which suggests that Netrin-1 signaling is unaffected, as DCC levels increase in *Netrin-1* mutants (Yung et al. 2015) (**Figure S4I-N**). Thus, the severe axon guidance phenotypes in *Dcc*^I372R/I372R^ mutant spinal cords do not seem to be due to an indirect effect on Netrin-1 activity.

### Changes in DCC diversity have similar effects on other Netrin-1-dependent neurons

Analysis of the retina, where Netrin-1 also acts both at short and long ranges, revealed the same range of effects that we observed in commissural axons. In the retina, Netrin-1 signals locally to guide DCC+ axons through the optic disc and out of the retina via the optic nerve (Deiner et al. 1997) (**Figure 5A**). *Dcc* mRNA is highly expressed in the retina during early embryonic development and decreases over time (**Figure 5B, S5A-C**). DCC protein is primarily localized to retinal ganglion cell (RGC) axons, which grow just under the RGC layer towards the optic disc (**Figure 5C,D**). Similar to *Ntn1*^-/-^ mutant embryos, optic nerves are hypoplastic in *Dcc*^-/-^ retinas, due in part to the re-routing of RGC axons to the subretinal space (Deiner et al. 1997; Vigouroux et al. 2020). *Draxin* mRNA was also detected in the embryonic retina and the resulting protein localized to RGC axons (**Figure S5 D-G**), raising the possibility that Draxin-DCC interactions could also play a role.

**Figure 5:**
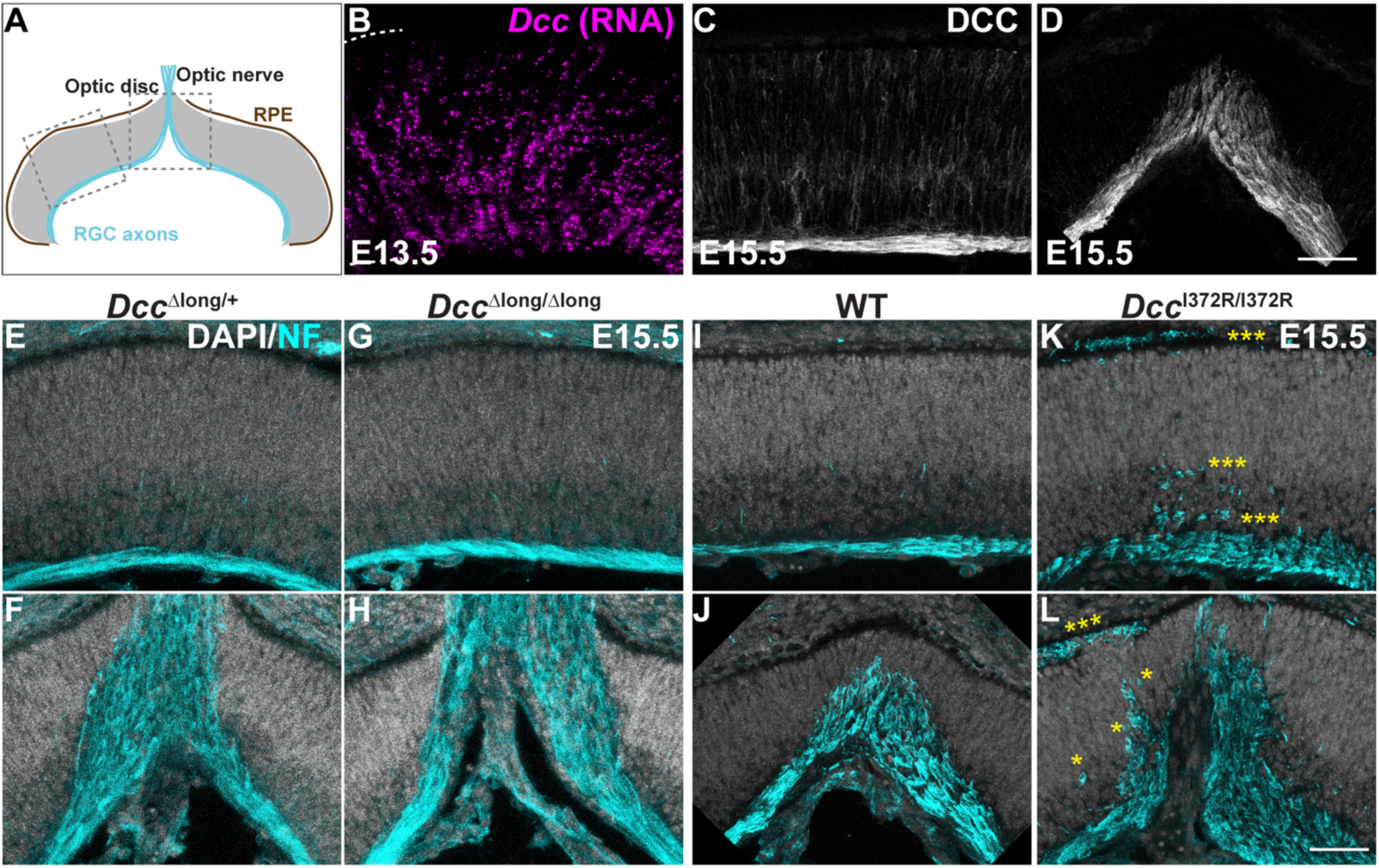
Some DCC ECD modules are required for RGC axons to properly exit the retina. **(A)** Representation of a retinal cross section. Squares indicate the ROIs in the following panels. **(B)** RNAscope *in situ* hybridization to *Dcc* mRNA on E13.5 retinal sections. **(C)** DCC protein localization on E15.5 retinal sections. **(D)** DCC protein localization on E15.5 retinal sections at the optic disc. **(E,F)** Representative image of DAPI and neurofilament (NF) staining on E15.5 retinal sections of *Dcc*^Δlong/+^ mice adjacent (E) and at the optic disc (F) (n=3 animals). **(G,H)** Representative image of DAPI and NF staining on E15.5 retinal sections of *Dcc*^Δlong/Δlong^ adjacent (G) and at the optic disc (H) (n=3 animals). **(I,J)** Representative image of DAPI and NF staining on E15.5 retinal sections of *wild type* (WT) mice adjacent (I) and at the optic disc (J) (n=3 animals). **(K,L)** Representative image of DAPI and NF staining on E15.5 retinal sections of *Dcc*^I372R/I372R^ mice adjacent (K) and at the optic disc (L) (n=3 animals). Scale bar: 50µm.

Similar to what was observed in the spinal cord, *Dcc*^I372/I372^ mutants exhibited stronger phenotypes in the retina than either *Dcc*^Δlong/Δlong^ or *Draxin*^-/-^ mutants. RGC axons exited the retina normally in *Dcc*^Δlong/Δlong^ animals (**Figure 5E-H**), indicating that DCC^short^ is sufficient for short range signaling here. Likewise, RGC axons did not grow ectopically into the retina in *Draxin*^-/-^ mutants (**Figure S5 H,I**). On the other hand, E15.5 *Dcc*^I372R/I372R^ mutants exhibited severely disrupted RGC guidance, much like what occurs in *Dcc*^-/-^ mutants. Neurofilament immunostaining revealed smooth growth of RGC axons through the neurofibrillary layer of the wildtype retina, but highly abnormal trajectories in the *DCC*^I372R/I372R^ mutant retina (**Figure 5I-L**). Specifically, there were fewer RGC axons in the neurofibrillary layer, with many extending ectopically into the retinal layers and subretinal space (asterisks in **Figure 5K**). In addition, *Dcc*^I372R/I372R^ mutant retinas showed aberrations at the optic disc, with more axons exiting through other areas (**Figure 5L**), similar to what occurs in *Dcc* null mutants. Thus, the I372R allele of DCC, which interferes with Draxin binding without disrupting Netrin-1 binding, again causes greater effects on axon guidance than the loss of Draxin. This suggests that this point mutation disrupts aspects of Netrin-1 signaling that are independent of Draxin.

To further test how each allele influences short range signaling by Netrin-1, we examined pontine motor neuron migration. Previous work revealed that Netrin-1 acts locally and in a non-gradient dependent fashion to confine migrating pontine motor neurons to the central nervous system (van Battum et al. 2025; Yung et al. 2018). Upon complete loss of Netrin-1 or DCC, pontine motor neuron precursors exit the hindbrain and migrate along the eighth cranial nerve towards the cochlea (Yung et al. 2018; Moreno-Bravo et al. 2018) (**Figure 6A**). In *Ntn1* null mutants, pontine motor neurons accumulate ectopically in the cochlea, but in *Dcc*^-/-^ mutants, the neurons do not migrate that far along the VIII^th^ nerve (Yung et al. 2018), suggesting that Netrin-1 influences this migration in multiple ways.

**Figure 6:**
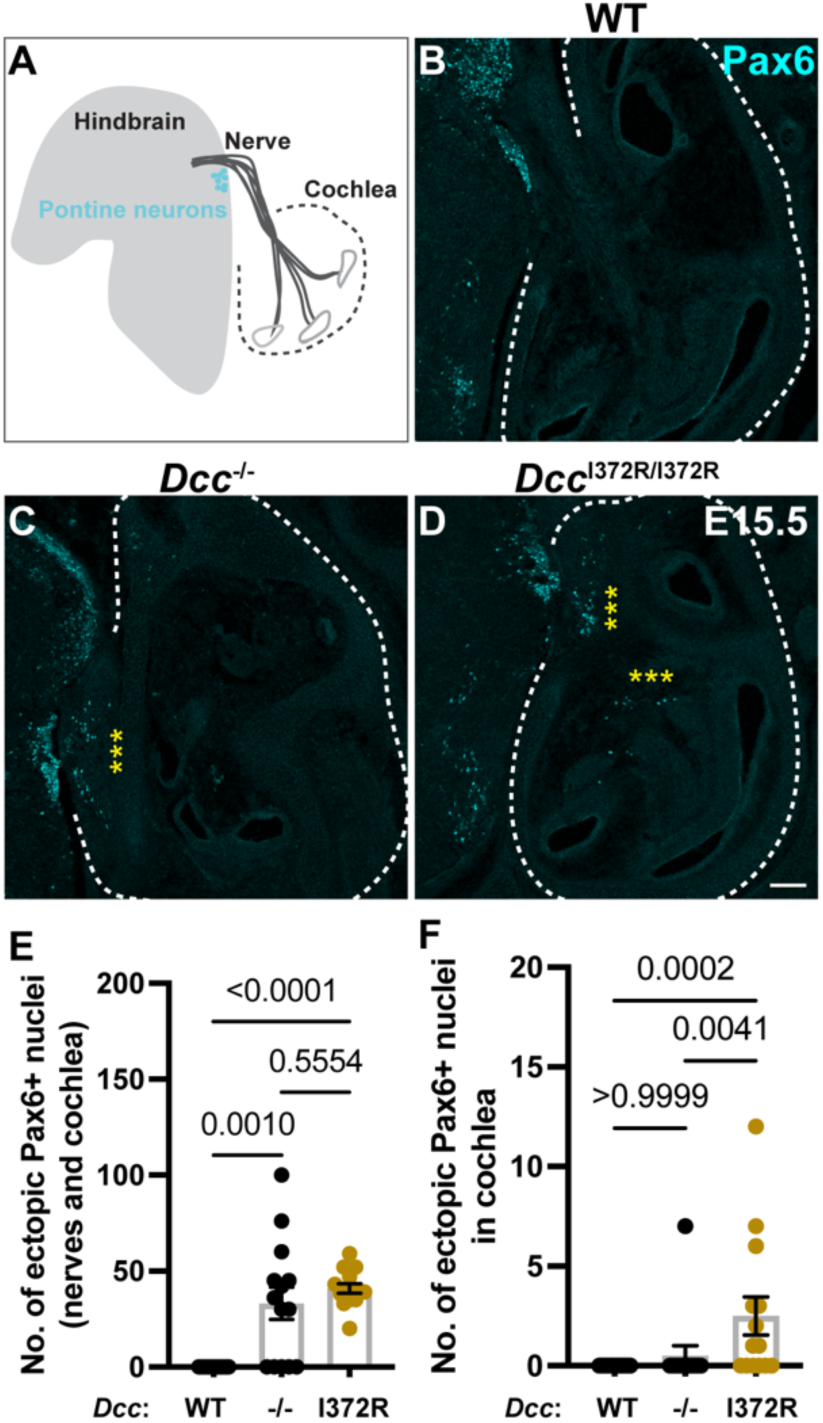
*Dcc*^I372R/I372R^ mutant mice exhibit pontine neuron mismigration into the developing cochlea. **(A)** Representation of the hindbrain containing pontine neurons (Pax6+) in close proximity to the cochlea. **(B,C,D)** Pax6 staining of WT (B), *Dcc*^-/-^ (C), and *Dcc*^I372R/I372R^ (D) cochleas in proximity to the hindbrain at E15.5. **(E)** Quantification of the number of ectopic Pax6+ nuclei found on nerves and cochlea. WT: 0.00 ± 0.00, n=19; *Dcc*^-/-^: 33.14 ± 8.42, n=14; *Dcc*^I372R/I372R^: 40.93 ± 2.57, n=14. Kruskal-Wallis test with multiple comparisons. **(F)** Quantification of the number of ectopic Pax6+ nuclei found in the cochlea only. WT: 0.00 ± 0.00, n=19; *Dcc*^-/-^: 0.50 ± 0.50, n=14; *Dcc*^I372R/I372R^: 2.50 ± 0.95, n=14. Kruskal-Wallis test with multiple comparisons. WT: 5 mice, *Dcc*^-/-^: 3 mice, *Dcc*^I327R/I372R^: 3 mice. Scale bar: 100µm.

Comparison of pontine neuron distribution in homozygous mutants for each of the DCC alleles suggests that the DCC^I372R^ mutation disrupts Netrin-1 signaling beyond its ability to bind to Draxin. We confirmed that DCC+ pontine neuron precursors migrate from the rhombic lip towards the ventral midline of the pons at E15.5 and that no Pax6+ pontine neurons are present in the VIII^th^ nerve or cochlea of wild-type animals (**Figure S6A,C,E** and **Figure 6B,E,F**). In *Dcc^-/-^* mutants, Pax6+ neurons mismigrate into the VIII^th^ nerve but do not reach the cochlea (**Figure 6C,E,F**). Unexpectedly, in *Dcc*^I372R/I372R^ mutants, Pax6+ pontine neurons accumulate not only in the VIII^th^ nerve but also in the cochlea (**Figure 6D,E,F**), similar to what occurs in *Ntn1* mutants (Yung et al. 2018). Thus, migration defects are stronger in *Dcc*^I372R/I372R^ mutants than in *Dcc*^-/-^. This observation suggests that the DCC^I372R^ molecule might disrupt additional modes of Netrin-1 signaling in a dominant negative fashion. No migration defects were observed in *Draxin^-/-^* or *Dcc*^Δlong/Δlong^ animals, consistent with the idea that these mutations primarily affect long-range Netrin-1 signaling (**Figure S6C-F**).

### The I372R mutation alters DCC clustering

The similarities between *Dcc*^I372R/I372R^ and *Dcc*^-/-^ mutants raised the possibility that a fundamental feature of DCC activation was lost upon changing a single residue in the ECD. Netrin-1 induces the formation of DCC clusters that are important for signaling (Bouchard et al. 2004). To determine how the I372R mutation might alter DCC signaling, we compared the distribution of total DCC and surface DCC by staining cultured commissural neurons from E13.5 WT and *Dcc*^I372R/I372R^ animals (**Figure 7A-D**). As expected, when prepared for total protein staining, NF (intracellular) was labeled successfully (**Figure 7A’,B’)** whereas no NF labeling was observed in neurons prepared for surface protein staining (**Figure 7C’,D’**). Total DCC was present at similar levels in WT and *Dcc*^I372R/I372R^ commissural axons (**Figure 7B**), consistent with tissue immunostaining and Western blot results (**Figure 1L, S2B**). Clusters of DCC were detected on the surface of WT commissural axons, even in the absence of exogenous Netrin-1 (**Figure 7C**). Indeed, DCC also clusters on cultured *Ntn1*^-/-^ commissural axons (**Figure S7A,B**). By contrast, although DCC^I372R^ protein trafficked to the surface, axons from *Dcc*^I372R/I372R^ animals did not form clear clusters (**Figure 7D**), confirmed by scoring the presence of clusters blind to genotype (**Figure 7E**). Although the I372R mutation disrupts Draxin-DCC binding (**Figure 1I-K**), commissural axons from *Draxin*^-/-^ mutants still formed DCC clusters on their surface and this distribution was unaffected by the addition of exogenous Draxin (**Figure S7C,D**). These results reinforce the conclusion that a change to the Draxin binding site also disrupts Draxin-independent features of DCC signaling. Additionally, DCC clustering on commissural axons was not altered in *Dcc*^Δlong/Δlong^ animals, where a greater portion (compared to *Dcc*^I372R/I372R^ animals) of the extracellular domain is absent (**Figure S7E-H**). Collectively, these findings demonstrate that specific residues in the DCC ECD are important for different downstream signaling events and hence reliable axon guidance.

**Figure 7:**
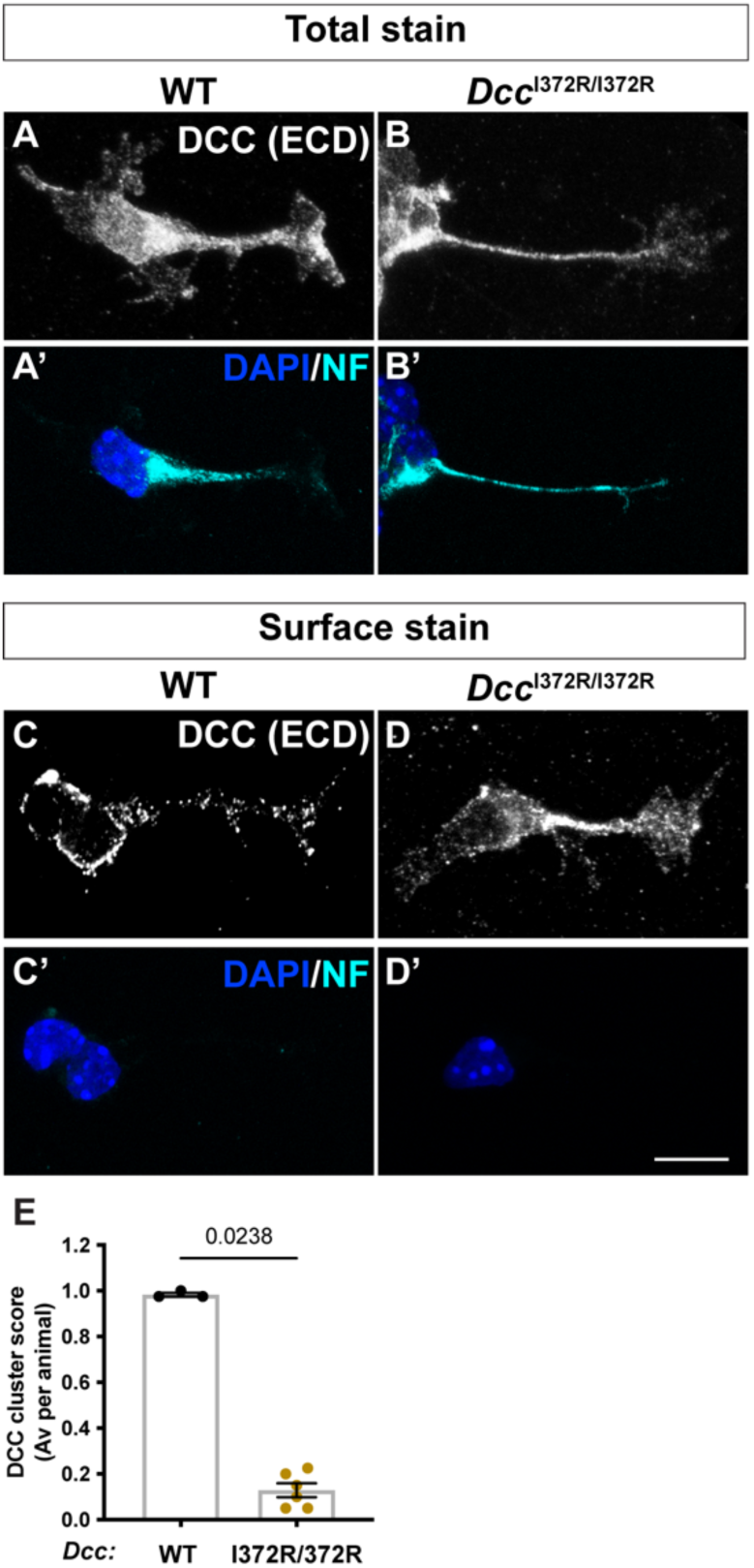
DCC does not cluster on the surface of *Dcc*^I372R/I372R^ mutant commissural axons. **(A,B)** DCC total staining of primary culture of WT (A) and *Dcc*^I372R/I372R^ (B) E13.5 commissural axons. A’ and B’ show the same culture as in A and B, respectively, stained for DAPI and NF. **(C,D)** DCC surface staining of primary culture of WT (C) and *Dcc*^I372R/I372R^ (D) E13.5 commissural axons. C’ and D’ show the same culture as in A and B, respectively, stained for DAPI and NF. **(E)** Quantification of DCC clusters on the surface of cultured commissural axons as a score where 1 indicates the presence of clusters. WT: 0.98 ± 0.01, n=3 animals; *Dcc*^I372R/I372R^: 0.13 ± 0.03, n=6 animals. Mann-Whitney test. Scale bar: 10µm.

## Discussion

The developing nervous system is patterned by cell surface proteins that can have fundamentally different effects depending on when and where they act. For example, molecular modularity in the Fat3 cytoplasmic domain leads to directed migration initially and synaptic localization subsequently (Avilés et al. 2022). Here, we show that molecular modularity in the extracellular domain of DCC ensures reliable axon guidance, with DCC^short^ playing a dominant role that is augmented by the long-range activity of DCC^long^. Further, alteration of a single residue sequentially far away from the Netrin-1 binding site strongly disrupted DCC activity. The presence of specific motifs that bind to different ligands and elicit distinct cellular responses diversifies the response of the growth cone as it traverses complex territories. As well as demonstrating an effective mechanism for reliable guidance, this functional dissection of the DCC ECD provides critical insight into the impact of human genetic mutations in DCC, which can have a range of clinical manifestations (Prato et al. 2024; Collins Hutchinson et al. 2024).

One site of developmentally regulated modularity in the DCC ECD is the linker between FN4 and FN5 domains. Structural observations suggested that this 20 amino acid long stretch could be crucial both for Netrin-1-induced oligomerization and Netrin-1’s long range functions (Finci et al. 2015, 2017, 2014; Priest et al. 2024). Here we show that the inclusion of this linker in DCC^long^ has subtle but significant effects on commissural axon guidance (**Figure 2H,L**), similar to what was described by others (Dailey-Krempel et al. 2023). The absence of stronger guidance defects suggests that DCC^short^ is able to mediate most of DCC’s effects on axon guidance, with additional compensation achieved by the ventral expansion of DCC^short^ in *Dcc*^Δlong/Δlong^ mutant spinal cords (**Figure 2D**). For instance, DCC^short^ on ventral neurons could enable local adhesive interactions that make up for the loss of long-range signaling activity. Indeed, loss of one *Ntn1* null allele from *Dcc*^Δlong/Δlong^ mutants revealed phenotypes previously linked to long range Ntn1 signaling (Wu et al. 2019), including invasion of the motor columns and a change in the shape of the trajectory towards the midline (**Figure 2I-M**). Thus, although DCC^short^ is able to take over many DCC^long^ functions, our data support the idea that DCC^long^ normally mediates long-range Netrin-1 dependent axon guidance.

DCC’s ability to bind to ligands other than Ntn1 creates a second source of molecular versatility that further enhances axon guidance. Among a large number of potential alternative ligands (Wojtowicz et al. 2020; Zhu et al. 2025), only Draxin has been investigated in detail. Draxin, also known as Neucrin, is highly expressed in the developing central nervous system (Miyake et al. 2009; Liu et al. 2018) and is critical for the formation of the zebrafish central nervous system, including the retina (Miyake et al. 2012). Consistent with its ability to bind DCC, multiple commissures are disrupted in the brains of *Draxin* mutant mice (Morcom et al. 2021; Ahmed and Shinmyo 2021; Islam et al. 2009; Shinmyo 2025), and we discovered additional defects in commissural axon guidance in the ventral spinal cord. However, Draxin/Neucrin also inhibits the canonical Wnt signaling pathway via interactions with LRP6 (Hutchins and Bronner 2018). Since Wnt signaling affects commissural axon guidance post midline crossing (Avilés and Stoeckli 2016; Avilés et al. 2013), it is possible that some Draxin-associated phenotypes are independent of its interactions with DCC. Consistent with this idea, Draxin phenotypes can be modulated by genetic background (Shinmyo 2025; Morcom et al. 2021).

Despite Draxin’s engagement with multiple signaling pathways across a range of tissues, *Draxin* null phenotypes are relatively mild, suggesting that like the insertion of the FN4-5 linker, the ability to bind Draxin has only nuanced effects on DCC’s activity. Whether Draxin and DCC^long^ act together or independently is unclear. Based on structural interactions, one possibility is that Draxin bridges DCC-Netrin-1 complexes across cells, perhaps enhancing DCC-Netrin-1 mediated adhesion and hence fasciculation. However, mutation of isoleucine 372 in the Draxin binding site essentially prevented DCC from steering axons across a range of tissues and created much stronger phenotypes than loss of Draxin. How this comes about at the molecular level is unclear. We considered the possibility that DCC^I372R^’s inability to bind to Draxin created an extra pool of Draxin that bound and sequestered Netrin-1, an antagonism that has been proposed (Gao et al. 2015). However, removal of Draxin did not rescue the *Dcc*^I372R/I372R^ phenotype (**Figure S4**). Since DCC^I372R^ protein still trafficked to the cell surface and could bind Netrin-1, an alternative explanation is that this mutation disrupted binding to ligands in addition to Draxin. In fact, the pontine neuron mismigration into the cochlea observed in *Dcc*^I372R/I372R^ mutants is not seen in *Dcc*^-/-^ embryos (**Figure 6**). Collectively, these observations suggest that the DCC ECD interacts with an unknown ligand through a binding site involving isoleucine 372 and that disruption of this binding frees the ligand to negatively affect axon and neuronal guidance.

Regardless of what ligands are involved, our results inform broader understanding of how DCC responds to cues in the environment. Previous studies showed that DCC receptors form clusters on axon surfaces in a Netrin-1-dependent manner (Matsumoto and Nagashima 2010; Bouchard et al. 2004). By contrast, we found that DCC clusters form on the surface of commissural neurons without application of exogenous Netrin-1 (**Figure 7C,S7A,G**) and also when no endogenous Netrin-1 is present (**Figure S7A,B**). These observations are consistent with the report that Netrin-1 promotes the growth of existing DCC nanoclusters, but not their nucleation (Uriot et al. 2024). Our results further highlight the importance of an alternative ligand, as the I372R mutation strongly reduces DCC’s capacity to cluster (**Figure 7D**) and no clustering deficits were observed in *Draxin* mutants (**Figure S7C,D**). The DCC ECD is sufficient for oligomerization, whereas the intracellular domain is necessary for recruitment to the cell surface (Gopal et al. 2016). One possibility is that the I372R mutation disrupts DCC oligomerization in response to an unknown cue, thereby preventing intracellular signaling needed to create clusters at specific sites on the cell surface. Therefore, our results support a model where a central-nervous-system-derived ligand promotes DCC cluster formation on the cell surface by interacting with DCC isoleucine 372 and thus influencing its ability to guide axons and neuronal cell bodies.

## Materials and Methods

### Animals

The novel *Dcc* mouse strains were generated by CRISPR/Cas9 zygote injection at the Genome Modification Facility at Harvard University. For *Dcc*^Δlong^, a gRNA targeting exon 17 (TCTGCTTTCTGCTTTGCTCA) produced a 25-nt deletion spanning intronic sequence and the 5’ end of exon 17. To generate *Dcc*^I372R^, we used a gRNA targeting exon 6 (CTAGCTTAGAAACGTAACCTCT) with an ssDNA donor containing the ATC→CGG mutation. All mutations were confirmed by sequencing the targeted loci and surrounding regions. The *Draxin*^-/-^ strain was provided by RIKEN (Accession CDB0484K)(Islam et al. 2009). *Dcc*^-/-^ and *Ntn1*^-/-^ mice were used and genotyped as previously described (Fazeli et al. 1997; Yung et al. 2015). Embryonic tissues were collected after confirming vaginal plugs the morning after mating. Mice were maintained on a 12/12 h light–dark cycle at 18–23 °C and 40–60% humidity. All procedures were approved by the Harvard Medical School IACUC. Genotyping was performed by real-time PCR (Transnetyx).

### RT-PCR

Total RNA was extracted from E11.5 heads using the RNeasy kit (Qiagen). For cDNA synthesis, 0.5 µg RNA was reverse-transcribed with random primers (Thermo Fisher Scientific, Cat#48190011), dNTPs, First-Strand buffer, DTT, RNaseOUT (Thermo Fisher Scientific, Cat#10777019), and SuperScript III (Thermo Fisher Scientific, Cat#18080044), incubated at 50 °C for 1 h and inactivated at 70 °C for 15 min. The resulting cDNA was amplified with PfuUltra II polymerase (Agilent, Cat# 600674) for 35 cycles (95 °C 20 s, 50 °C 20 s, 72 °C 30 s). Isoform expression was assessed using primers spanning exon junctions. Long (237 bp) and short (177 bp) isoforms were amplified with EA226/EA229. The short isoform (166 bp) was detected with EA228/EA229, and the long (228 bp) with EA227/EA229. PCR products were run on 3% agarose gels.

### Protein interaction in Cos7 cells

Cos7 cells were cultured in DMEM + 10% FBS at 37 °C with 5% CO_2_. DCC^WT^ and DCC^I372R^ plasmids were transfected using Lipofectamine 2000 (Thermo Fisher Scientific, Cat#11668019). After 30 hours, cells were incubated for 90 min at 4 °C with recombinant Netrin-1 (R&D systems, 1109-N1-025) or Draxin (R&D systems, 6148-DR-025) (200 ng/mL) in protein-binding buffer (20mM HEPES, 0.2% BSA, 5mM CaCl_2_, 1mM MgCl_2_, 2µg/mL Heparin in HBSS). Following washes, cells were stained for DCC, Netrin-1, and Draxin. Mean fluorescence was quantified per image and normalized to background.

### Western blot

E13.5 heads were lysed in N-PER buffer (Thermo Scientific 87792), and clarified lysates were collected after centrifugation (16,000 g, 30 min, 4°C). Samples were denatured at 95°C for 10 min and run on 4–12% Bis-Tris gels (Bio-Rad Cat#3450124) in MES buffer (Bio-Rad Cat#1610789). Proteins were transferred (1 h, 75 V) to Immobilon-P membranes (0.45µm, Sigma-Millipore) and blocked in 5% milk. Membranes were incubated overnight at 4°C with mouse anti-DCC (1:200) and rabbit anti-β-actin (1:1,000). After TBS-T washes, HRP-conjugated secondary antibodies (1:2,000) were applied for 1-2 h at room temperature. Signals were developed using Clarity ECL substrate (Bio-Rad, Cat#1705061).

### Immunohistochemistry, and *In situ* hybridization (RNAscope and BaseScope)

Animals at the required embryonic or postnatal ages were euthanized by CO_2_ inhalation and cervical dislocation. Embryonic tissues were collected at E11.5 (spinal cord) and E15.5 (heads for retinas and hindbrain/cochlea). For postnatal retinas, extraocular tissues were removed, keeping the eyecups. All samples were fixed in 4% PFA (PFA, EMS Cat#15710) for 30 min, washed in PBS, cryoprotected in 30% sucrose for 2 h at 4°C, incubated in NEG-50 (VWR, Cat#84000-154) overnight, and embedded by freezing. Cryosections (20 µm) of spinal cord, heads, or postnatal eyes were mounted on Superfrost Plus slides (VWR, Cat#48311-703). Spinal cord sections were taken from the lumbar region. For immunohistochemistry, NEG-50 was removed in PBS, and sections were blocked/permeabilized in 5% normal donkey serum (NDS, Jackson ImmunoResearch Cat#017-000-121) with 0.5% Triton-X for 1-2 h. Samples were incubated with primary antibodies overnight at 4°C, washed, then incubated with fluorescent secondary antibodies in 5% NDS with 0.02% Triton-X for 1.5–2 h. After final washes, slides were mounted in DAPI-Fluoromount-G (SouthernBiotech Cat#0100-20). Antibody details are provided in Table S1. RNAscope fluorescent *in situ* hybridization was performed using the RNAscope Multiplex v2 kit (ACD, Cat#323120), following manufacturer instructions. Tissue was collected as for immunohistochemistry but under RNase-free conditions. Post-fixation in 4% PFA for 15 min, hydrogen peroxide for 10 min, and Protease III for 10 min at 40°C before probe incubation (probes listed in Table S1). After RNAscope, sections were rinsed, re-blocked in 5% NDS/0.5% Triton X-100, and processed for standard immunohistochemistry. Isoform-specific detection used the BaseScope RED kit (ACD, Cat#322900), with the same pretreatments (PFA 15 min, hydrogen peroxide 10 min, Protease III 10 min at 40°C). BaseScope probes (Table S1) included BA-Mm-Dcc-E16E17 for the long isoform (exon 16–17 junction) and BA-Mm-Dcc-tvX1-E16E17 for the short isoform. Tissue sections were imaged with a Zeiss LSM800 or Leica SP8 confocal microscopes. The entire sections were imaged in consecutive z-slices separated by 1 µm using a 20x, or 40x oil objective. The z stacks were then projected at maximum fluorescence intensity using Fiji/ImageJ. For quantifications, images were randomly numbered for blind quantification, and all analyses were performed under identical technical conditions, comparing control and experimental samples within the same experiment. Robo3⁺ axon endings in motor column, dorsal, and ventral regions were visually counted per spinal cord section. Commissure density was measured in a 73.86 × 37.5 µm midline area using ImageJ after thresholding (**Figure S1)**. The distance from the main Robo3⁺ bundle to the midline was also measured in ImageJ. Axon fasciculation was assessed by calculating the ratio a/b, where “a” is the distance from the lateral spinal cord to commissural axons and “b” is the distance to the midline (**Figure 3A**); larger ratios indicate defasciculation. Pax6⁺ nuclei in nerves and cochlea were counted visually per section.

### Open book and whole mount immunostaining of spinal cords

Spinal cords from E11.5 embryos were dissected in PBS, pinned down as open books, and fixed in 4% PFA for 30 min. After washes, tissues were blocked overnight in PBS with 0.2% gelatin and 0.5% Triton X-100. Anti-Robo3 antibody was applied in blocking buffer and incubated at 37°C for 5 days. Following washes, Alexa 568 secondary antibody was added overnight at 37°C. Spinal cords were washed again and mounted in DAPI-Fluoromount.

### *In vitro* commissural neuron culture

Embryonic commissural neuron cultures were prepared following Langlois et al. (2010) (Langlois et al. 2010). E13.5 embryos were dissected in L-15 on ice; spinal cords were collected in 10% FBS/L-15 lying as flat as possible (open book), the dorsal fifth was removed, and the tissue was kept in 10% FBS/L-15. After washing in cold HBSS, strips were digested in 0.15% trypsin for 7 min at 37°C, then treated with DNase and MgSO_4_ and centrifuged. Pellets were resuspended in Neurobasal + 10% FBS + L-Glutamine and triturated. Cells were plated on Poly-L-Lysine and, after 16–18 h, switched to Neurobasal + B27 + L-Glutamine. For surface staining, cultures were cooled, medium removed, and primary antibodies added in growth medium for 1 h at 4°C, followed by fixation in 4% PFA. For total staining, cells were fixed first, then permeabilized in 0.1% Triton X-100/PBS and incubated with primary antibodies overnight at 4°C. After PBS washes, fluorescent secondaries were applied for 1.5 h, and cells were mounted in DAPI-Fluoromount. Cluster scores were assigned per cell: 1 (with clusters), 0.5 (partial presence of clusters), or 0 (no cluster).

### Statistics and data availability

Statistical analyses were performed in Prism10. Two-tailed t tests were used for Gaussian data and Mann–Whitney tests for non-Gaussian data. For comparisons among more than two groups, one-way ANOVA with Dunnett’s test was applied. Sample size (n), mean, SEM, and p values are reported in the figure legends. Source data for graphs accompany the paper, and all unique materials and original images are available upon request.

## Acknowledgements

We thank members of the Goodrich, Segal, and Rao labs at Harvard Medical School for their insightful comments; RIKEN for sharing the Draxin null mouse strains; NIH grant R21 NS113562 (LVG); and a Leonard and Isabelle Goldenson Fellowship, Alice and Joseph Brooks Fund Fellowship, ANID Becas Chile Postdoctoral Fellowship, IBRO Rising stars, eLife Ben Barres Spotlight, and ANID Fondecyt de Iniciación Código 11241253 (ECA).

## Author Contributions

Conceptualization, ECA, LVG, AY, RM and JHW; methodology, ZD, AY; investigation, ECA; writing, ECA, RM, LVG; funding acquisition, ECA, LVG; supervision, ECA, LVG.

