## Supplemental material for "Modularity in the DCC extracellular domain elicits distinct effects on axon guidance"

**Avilés_Fig S1**

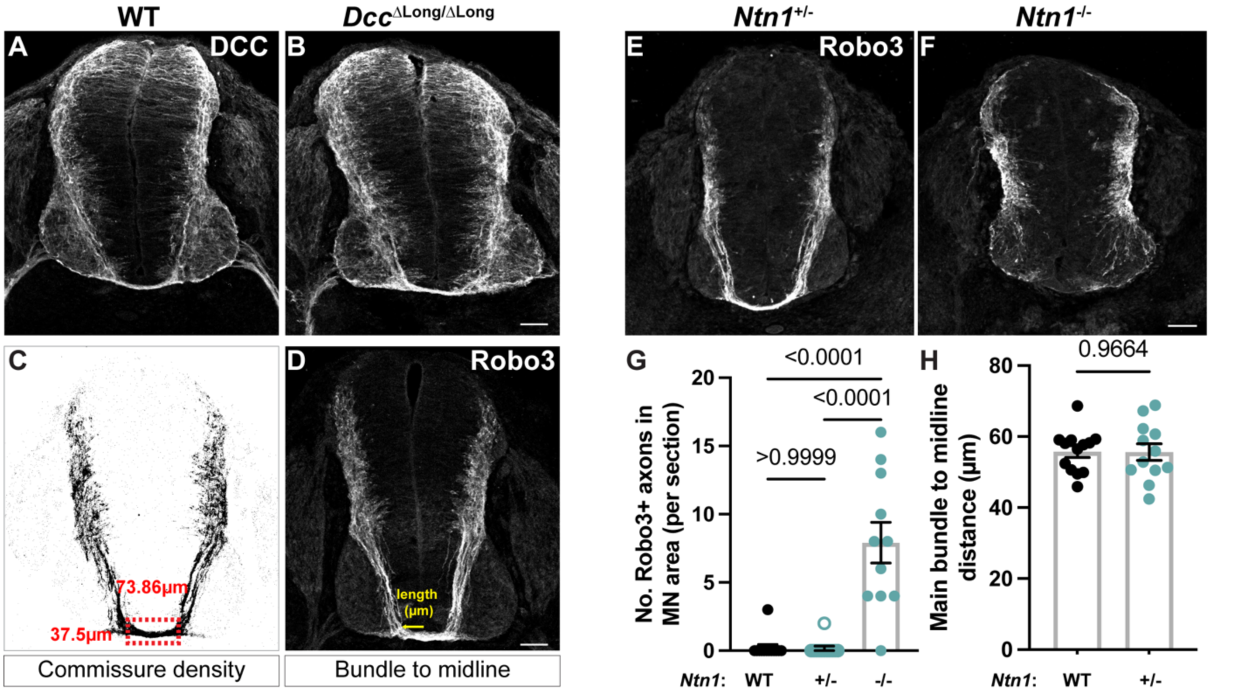

**<u>Figure S1</u>: $Dcc^{\Delta long/\Delta long}$ mice produce DCC that locates properly in the spinal cord. (A,B)** DCC staining of wild type (A) and $Dcc^{\Delta long/\Delta long}$ mutant (B) spinal cords. **(C)** Binarized image used to measure the integrated density on a region of interest, corresponding to 73.86 µm x 37.5 µm of area of the commissure. **(D)** Robo3 immunostaining of a spinal cord showing the measurement of the distance between the main axon bundle and the midline. **(E,F)** Robo3 immunostaining of *Ntn1*$^{+/-}$ (D) and *Ntn1*$^{-/-}$ (F) E11.5 spinal cords. **(G)** Quantification of the number of Robo3+ axons in the motor neuron area. WT: 0.23 ± 0.23, n=13; *Ntn1*$^{+/-}$: 0.17 ± 0.17, n=12; *Ntn1*$^{-/-}$: 7.90 ± 1.50, n=11. Kruskal-Wallis test. **(H)** Quantification of the distance between main bundle to the midline (µm). WT: 55.75 ± 1.65, n=13; *Ntn1*$^{+/-}$: 55.63 ± 2.35. *t* test. WT: 3 mice, *Ntn1*$^{+/-}$: 3 mice, *Ntn1*$^{-/-}$: 3 mice. Scale bars: 50µm.

Avilés_Fig S2

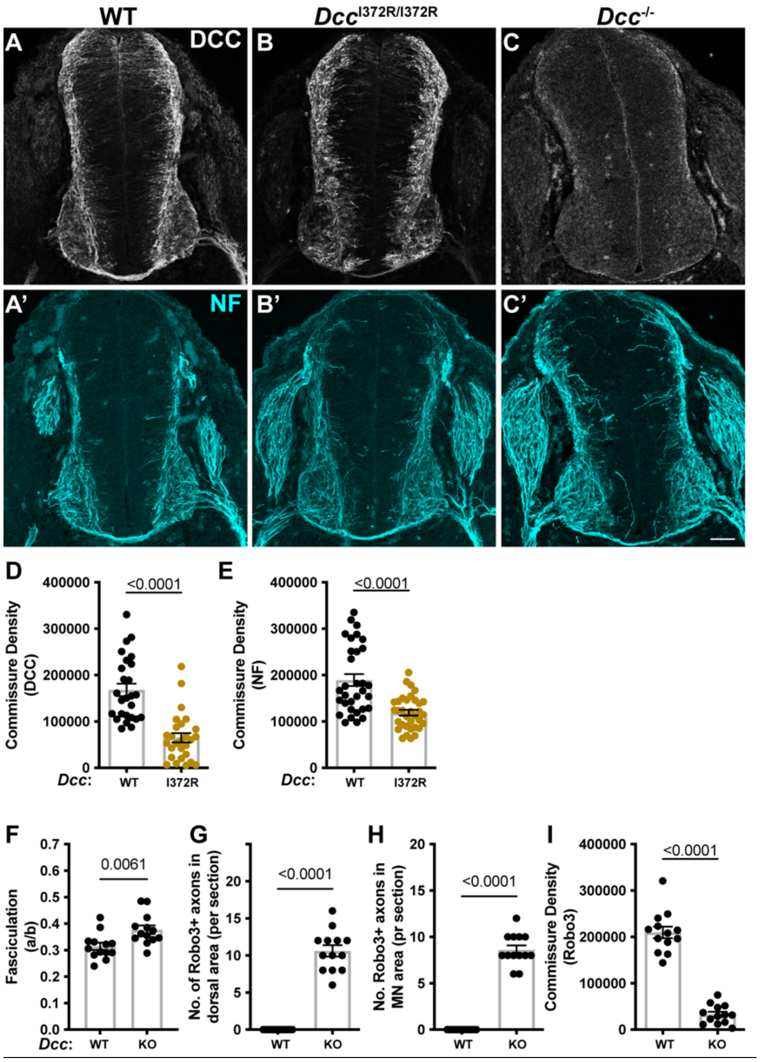

**Figure S2**: ***Dcc*$^{I372R/I372R}$ mutant spinal cords resemble the *Dcc* null phenotype. (A)** Immunostaining for DCC on a WT spinal cord cross section at E11.5. A' shows the same section stained for NF. **(B)** Immunostaining for DCC on a *Dcc*$^{I372R/I372R}$ spinal cord cross section at E11.5. B' shows the same section stained for NF. **(C)** Immunostaining for DCC on a *Dcc*$^{-/-}$ spinal cord cross section at E11.5. C' shows the same section stained for NF. **(D)** Quantification of the commissure density by DCC staining. WT: 168213 ± 13392, n=26; *Dcc*$^{I372R/I372R}$: 64759 ± 9950, n=27; Mann-Whitney test. **(E)** Quantification of the commissure density by NF staining. WT: 189285 ± 12938, n=32; *Dcc*$^{I372R/I372R}$: 118895 ± 6166, n=34; *t* test. WT: 8 mice, *Dcc*$^{I372R/I372R}$: 8 mice (D,E). **(F)** Quantification of the fasciculation parameter (a/b). WT: 0.31 ± 0.01, n=13; *Dcc*$^{-/-}$: 0.38 ± 0.02, n=13; *t* test. **(G)** Quantification of the number of Robo3+ axons in the dorsal area. WT: 0.00 ± 0.00, n=13; *Dcc*$^{-/-}$: 10.62 ± 0.76, n=13; Mann-Whitney test. **(H)** Quantification of the number of Robo3+ axons in the MN area. WT: 0.00 ± 0.00, n=13; *Dcc*$^{-/-}$: 8.62 ± 0.47, n=13; Mann-Whitney test. **(I)** Quantification of the commissure density by Robo3 staining. WT: 209755 ± 12487, n=13; *Dcc*$^{-/-}$: 32816 ± 5746, n=13; *t* test. WT: 4 mice, *Dcc*$^{-/-}$: 4 mice (F-I). Scale bar: 50μm.

Avilés_Fig S3

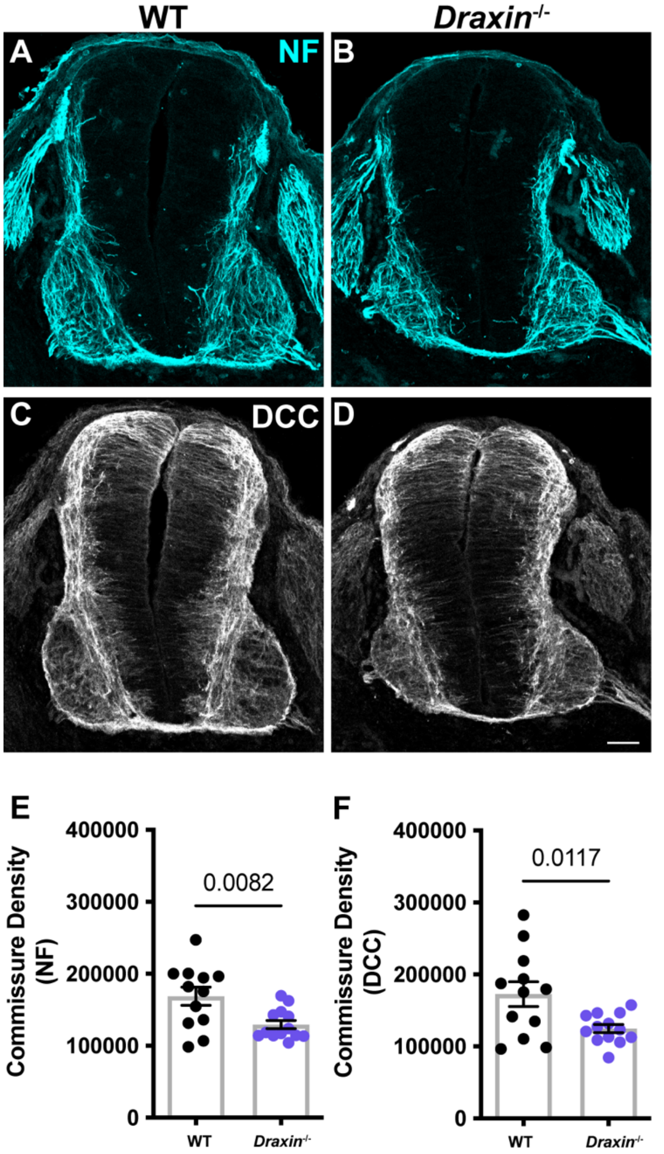

**Figure S3: Neurofilament and DCC immunostaining show a reduction of the commissure in *Draxin*$^{-/-}$ mutants. (A,B)** Neurofilament (NF) immunostaining of wild type (A) and *Draxin*$^{-/-}$ mutant (B) E15.5 spinal cords. **(C,D)** DCC immunostaining of wild type (C) and *Draxin*$^{-/-}$ mutant (D) E15.5 spinal cords. **(E)** Quantification of the commissure density by NF staining. WT: 168707 ± 12673, n=12; *Draxin*$^{-/-}$: 129442 ± 5766, n=13. Mann-Whitney test. **(F)** Quantification of the commissure density by DCC staining. WT: 172769 ± 17209, n=12; *Draxin*$^{-/-}$: 124791 ± 5643, n=13. *t* test. WT: 3 mice, *Draxin*$^{-/-}$: 3 mice. Scale bar: 50μm.

Avilés_Fig S4

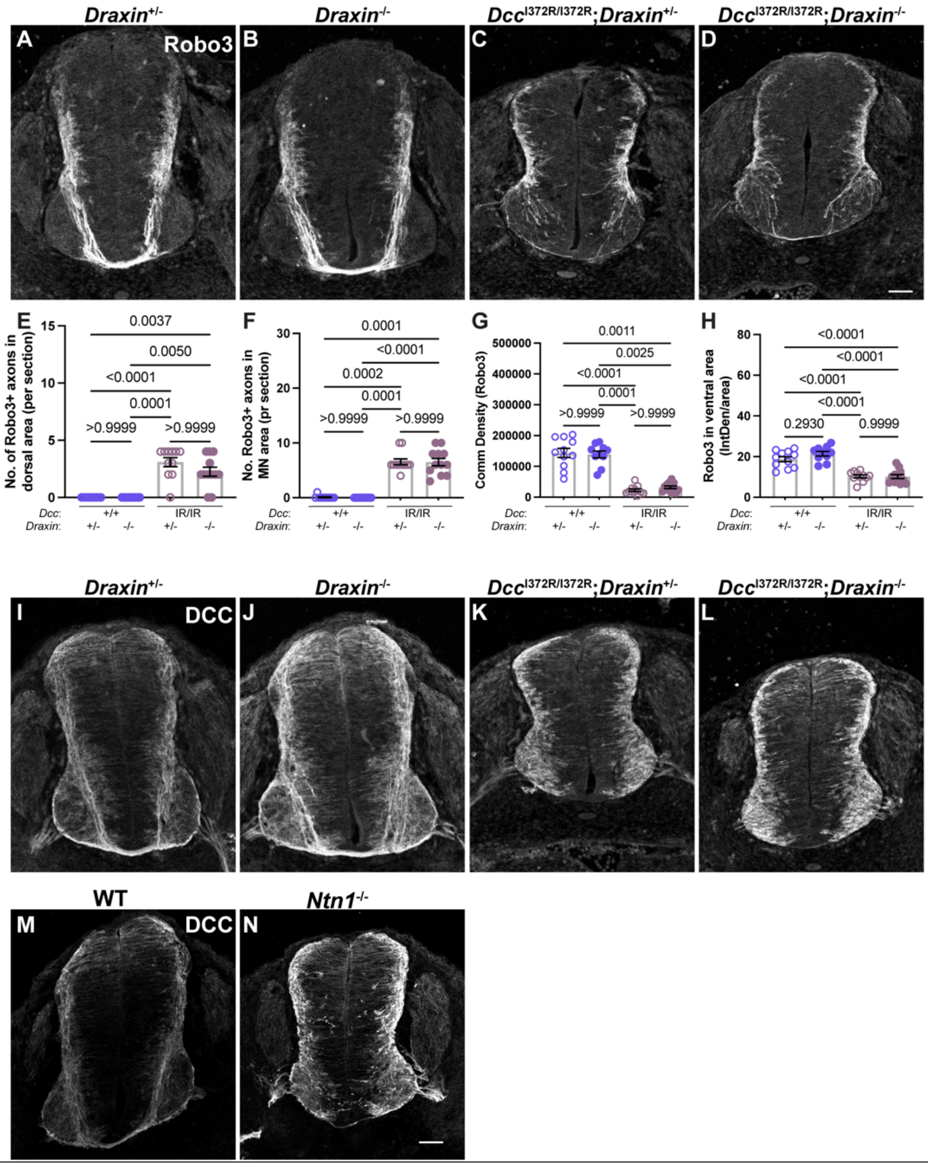

**Figure S4: Loss of *Draxin* does not counteract the *Dcc*$^{I372R/I372R}$ phenotype. (A-D)** Robo3 imunostaining of *Draxin*$^{+/-}$ (A), *Draxin*$^{-/-}$ (B), *Dcc*$^{I372R/I372R}$;*Draxin*$^{+/-}$ (C), and *Dcc*$^{I372R/I372R}$;*Draxin*$^{-/-}$ (D) E11.5 spinal cord cross sections. **(E)** Quantification of the number of Robo3+ axons in the dorsal area. *Draxin*$^{+/-}$: 0.00 ± 0.00, n=11; *Draxin*$^{-/-}$: 0.00 ± 0.00, n=10; *Dcc*$^{I372R/I37R}$; *Draxin*$^{+/-}$: 3.09 ± 0.39, n=11; *Dcc*$^{I372R/I37R}$; *Draxin*$^{-/-}$: 2.25 ± 0.39, n=12. Kruskal-Wallis test, with multiple comparisons. **(F)** Quantification of the number of Robo3+ axons in the MN area. *Draxin*$^{+/-}$: 0.09 ± 0.09, n=11; *Draxin*$^{-/-}$: 0.00 ± 0.00, n=10; *Dcc*$^{I372R/I37R}$; *Draxin*$^{+/-}$: 6.54 ± 0.54, n=11; *Dcc*$^{I372R/I37R}$; *Draxin*$^{-/-}$: 6.50 ± 0.67, n=12. Kruskal-Wallis test, with multiple comparisons. **(G)** Quantification of the commissure density by Robo3 staining. *Draxin*$^{+/-}$: 143741 ± 14634, n=11; *Draxin*$^{-/-}$: 138791 ± 11044, n=10; *Dcc*$^{I372R/I37R}$; *Draxin*$^{+/-}$: 22615 ± 4233, n=11; *Dcc*$^{I372R/I37R}$; *Draxin*$^{-/-}$: 32293 ± 4296, n=12. Kruskal-Wallis test, with multiple comparisons. **(H)** Quantification of Robo3+ axons in ventral area. *Draxin*$^{+/-}$: 18.75 ± 1.24, n=11; *Draxin*$^{-/-}$: 21.42 ± 1.15, n=10; *Dcc*$^{I372R/I37R}$; *Draxin*$^{+/-}$: 10.29 ± 0.73, n=11; *Dcc*$^{I372R/I37R}$; *Draxin*$^{-/-}$: 10.19 ± 0.96, n=12. One-way ANOVA test, with multiple comparisons. 3 mice for each condition. Scale bar: 50μm. **(I-L)** DCC imunostaining of *Draxin*$^{+/-}$ (I), *Draxin*$^{-/-}$ (J), *Dcc*$^{I372R/I372R}$;*Draxin*$^{+/-}$ (K), and *Dcc*$^{I372R/I372R}$;*Draxin*$^{-/-}$ (L) E11.5 spinal cord cross sections. **(M,N)** DCC immunostaining of WT (E) and *Ntn1*$^{-/-}$ E11.5 spinal cord cross sections. Scale bar: 50μm.

Avilés_Fig S5

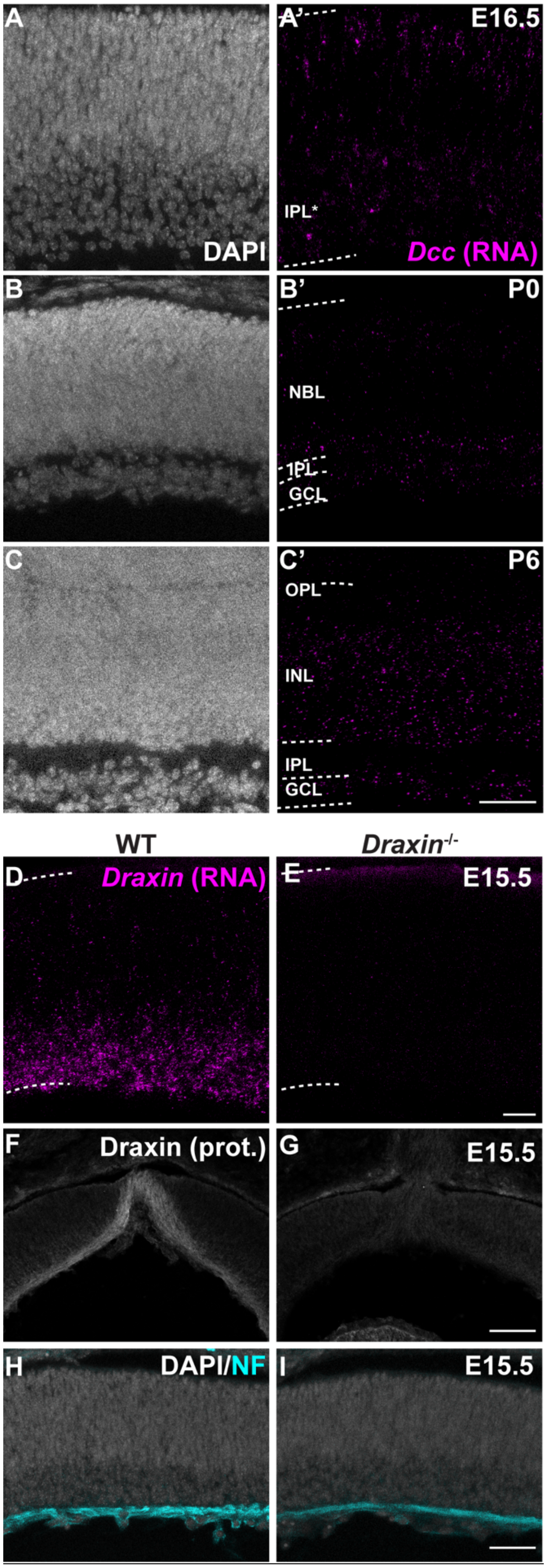

**<u>Figure S5</u>: *Dcc* RNA is expressed in the developing retina. (A)** RNAscope *in situ* hybridization to *Dcc* mRNA on E16.5 retinal sections. **(B)** RNAscope *in situ* hybridization to *Dcc* mRNA on P0 retinal sections. **(C)** RNAscope *in situ* hybridization to *Dcc* mRNA on P6 retinal sections. **(D)** RNAscope *in situ* hybridization to *Draxin* mRNA on WT retinal sections at E15.5. **(E)** RNAscope *in situ* hybridization to *Draxin* mRNA on *Draxin*$^{-/-}$ retinal sections at E15.5. **(F)** Immunostaining for Draxin on WT E15.5 retina. **(G)** Immunostaining for Draxin on *Draxin*$^{-/-}$ E15.5 retina. **(H)** Immunostaining for NF on WT E15.5 retina. **(I)** Immunostaining for NF on *Draxin*$^{-/-}$ E15.5 retina. Scale bars: 50μm (C); 20μm (I).

**Avilés_Fig S6**

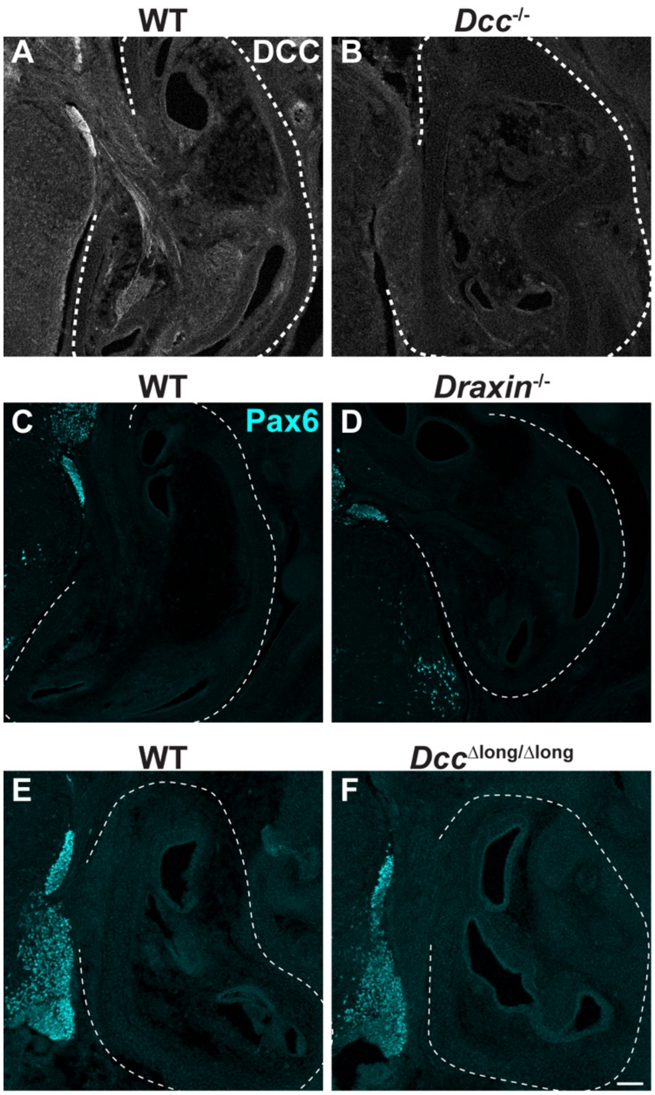

<u>Figure S6</u>: ***Draxin*$^{-/-}$ and *Dcc*$^{\Delta long/\Delta long}$ mutants do not show Pax6+ neuron mismigration to the cochlea.** **(A)** DCC immunostaining on WT cross sections of E15.5 cochlea. **(B)** DCC immunostaining on *Dcc*$^{-/-}$ cross sections of E15.5 cochlea. **(C)** Pax6 immunostaining on WT cross sections of E15.5 cochlea (littermates of D). **(D)** Pax6 immunostaining on *Draxin*$^{-/-}$ cross sections of E15.5 cochlea. **(E)** Pax6 immunostaining on WT cross sections of E15.5 cochlea (littermates of F). **(F)** Pax6 immunostaining on *Dcc*$^{\Delta long/\Delta long}$ cross sections of E15.5 cochlea. Scale bar: 100µm.

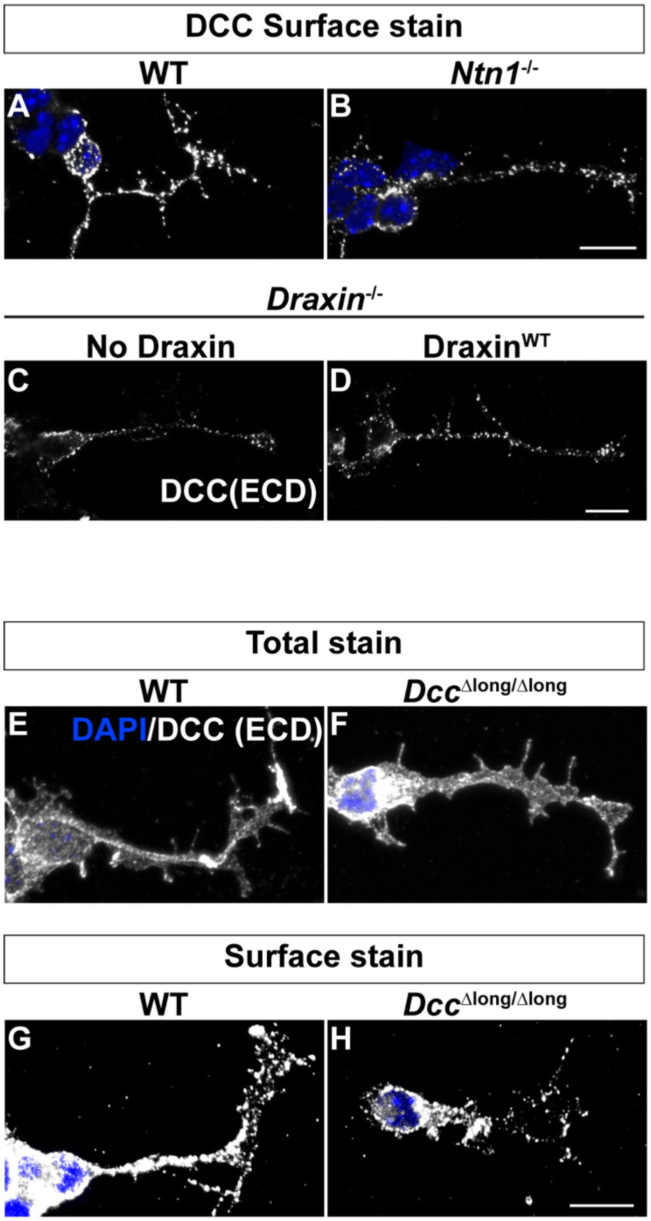

**<u>Figure S7</u>: DCC protein clusters on the surface of *Netrin*$^{-/-}$, *Draxin*$^{-/-}$, and *Dcc*$^{\Delta long/\Delta long}$ mutant commissural axons. (A)** DCC surface staining of primary culture of WT E13.5 commissural axons. **(B)** DCC surface staining of primary culture of *Netrin*$^{-/-}$ E13.5 commissural axons. **(C)** DCC surface staining of primary culture of *Draxin*$^{-/-}$ E13.5 commissural axons with no Draxin added to the culture. **(D)** DCC surface staining of primary culture of *Draxin*$^{-/-}$ E13.5 commissural axons with Draxin wild type added to the culture. **(E)** DCC total staining of primary culture of WT E13.5 commissural axons. **(F)** DCC total staining of primary culture of *Dcc*$^{\Delta long/\Delta long}$ E13.5 commissural axons. **(G)** DCC surface staining of primary culture of WT E13.5 commissural axons. **(H)** DCC surface staining of primary culture of *Dcc*$^{\Delta long/\Delta long}$ E13.5 commissural axons. Scale bar: 10μm.

**Table S1: Key resource table**

| Reagent or resource | Source | Identifier |
|---|---|---|
| **Antibodies** | | |
| Goat anti DCC | R&D systems | Cat# AF844;<br>RRID: AB_2089765 |
| Mouse anti DCC | Santa Cruz | Cat# sc-515834 |
| Rabbit anti β-actin | Raybiotech | Cat# 168-10656;<br>RRID: RRID:AB_2885189 |
| Goat anti Netrin-1 | R&D systems | Cat# AF1109;<br>RRID: RRID:AB_2298775 |
| Sheep anti Draxin | R&D systems | Cat# AF6149;<br>RRID: RRID:AB_10640005 |
| Goat anti Robo3 | R&D systems | Cat# AF3076;<br>RRID: RRID:AB_2181865 |
| Mouse anti Neurofilament | Developmental studies hybridoma bank | Cat# 2H3;<br>RRID: AB_531793 |
| Rabbit anti Pax6 | Biolegend | Cat# 901301;<br>RRID: AB_2565003 |
| Donkey anti goat, Alexa Fluor® 568 | Thermo Fisher Scientific | Cat#A11057;<br>RRID:AB_142581 |
| Donkey anti mouse, Alexa Fluor® 488 | Abcam | Cat#ab150105;<br>RRID:AB_2732856 |
| Donkey anti mouse, Alexa Fluor® 568 | Thermo Fisher Scientific | Cat#A10037;<br>RRID:AB_2534013 |
| Goat anti mouse, Alexa Fluor® 647 | Thermo Fisher Scientific | Cat#A-21235;<br>RRID:AB_2535804 |
| Donkey anti rabbit, Alexa Fluor® 488 | Thermo Fisher Scientific | Cat#A21206;<br>RRID:AB_2535792 |

| | | |
|---|---|---|
| Donkey anti rabbit, Alexa Fluor® 568 | Thermo Fisher Scientific | Cat#A10042; RRID:AB_2534017 |
| Donkey anti rabbit, Alexa Fluor® 647 | Thermo Fisher Scientific | Cat#A31573; RRID:AB_2536183 |
| Donkey anti sheep, Alexa Fluor® 568 | Thermo Fisher Scientific | Cat#A-21099; RRID:AB_2535753 |
| Goat anti mouse – HRP | BioRad | Cat# 170-6516; RRID:AB_11125547 |
| Goat anti rabbit – HRP | BioRad | Cat# 170-6515; RRID: AB_11125142 |
| **Commercial kits and assays** | | |
| RNeasy kit | Qiagen | Cat#74104 |
| RNAscope® Multiplex Fluorescent Reagent Kit v2 | ACD | Cat#323100 |
| BaseScope™ Reagent Kit v2- RED | ACD | Cat#323910 |
| **Experimental models: Organisms/Strains** | | |
| Mouse: $Dcc^{I372R}$ | This paper | N/A |
| Mouse: $Dcc^{\Delta long}$ | This paper | N/A |
| Mouse: $Ntn1^{-/-}$ | (Yung et al. 2015) | MGI:5888901 |
| Mouse: $Dcc^{-/-}$ | (Fazeli et al. 1997) | MGI:1934924 |
| Mouse: $Draxin^{-/-}$ | (Islam et al. 2009) | MGI:3833678 |
| ***In situ* hybridization Probes** | | |
| RNAscope™ Probe- Mm-Dcc | ACD | Cat# 427491 |

| | | |
|---|---|---|
| RNAscope® Probe- Mm-Draxin-C2 | ACD | Cat# 494231-C2 |
| BaseScope™ Probe- BA-Mm-Dcc-E16E17 | ACD | Cat# 708051 |
| BaseScope™ Probe- BA-Mm-Dcc-tvX1-E16E17 | ACD | Cat# 708061 |
| **Recombinant DNA** | | |
| DCC-WT plasmid | This paper | N/A |
| DCC-I372R plasmid | This paper | N/A |
| **PCR Primers** | | |
| EA226 | This paper | GCCACCACTCGTTCCATAAC |
| EA227 | This paper | GTTCCATAACAGATCCCACTGA |
| EA228 | This paper | TCCATAACAG\|ATGTCTCCACCC |
| EA229 | This paper | CAGAAAAGCTGGTCCTCCAC |
